# Kinbiont: From time series to ecological and evolutionary responses in microbial systems

**DOI:** 10.1101/2024.09.09.611847

**Authors:** Fabrizio Angaroni, Alberto Peruzzi, Edgar Z. Alvarenga, Fernanda Pinheiro

## Abstract

Microbial behavior is quantitatively characterized by observables inferred from kinetics experiments. Growth rate and biomass yield, for example, are used to map response patterns across different conditions including antibiotic growth inhibition and yield dependence on substrate. As microbial kinetics datasets grow, there is immense potential to advance our understanding of ecological and evolutionary processes. But how can we turn these data into actionable insights about microbial responses? Here we introduce Kinbiont – an ecosystem of numerical methods integrating advanced ordinary differential equation solvers, non-linear optimization, signal processing, and interpretable machine learning algorithms. Kinbiont offers a model-based data analysis pipeline covering all aspects of microbial kinetics, from pre-processing to result interpretation. We demonstrate Kinbiont’s performance using synthetic and real datasets, including bacterial growth, diauxic curves, phage-bacteria co-cultures, and ecotoxicological responses. Kinbiont can aid biological discovery through data-driven generation of hypotheses that can be tested in targeted experiments.

## Main

In the early 1940s, after meticulously gathering microbial growth data, Jacques Monod observed that the specific growth rate of bacteria exhibited a clear pattern in response to changes in substrate concentration, mirroring enzyme kinetics [1]. This observation led him to formulate a mathematical model that transformed our understanding of microbial growth. What Monod did is similar in kind to the process of symbolic regression: he discovered a symbolic expression that matched the data.

Today, microbial behavior in response to ecological and evolutionary perturbations is at the heart of modern scientific and medical challenges. The need to understand and predict the conditions under which bacteria become resistant to antibiotics, ecosystems remain resilient to interventions, and bioproduction reaches optimal efficiency has led to a surge in systematic experiments generating massive datasets. However, transforming these data into intervention strategies remains a significant challenge. It requires models that can generate testable hypotheses about microbial behavior in unobserved conditions [2, 3]. Developing these models involves establishing quantitative relationships between biological observables (e.g., growth rate, lag time, or yield) and ecological variables (e.g., nutrients, stressors, or inter-organismal interactions) or evolutionary factors (e.g., mutations) – what we refer to as ecological or evolutionary responses. Ecological responses map phenotypic patterns of an organism under challenges, such as antibiotic or nutrient stress; evolutionary responses map phenotypic differences among genetic variants in a given ecological setting. These phenotypes must first be inferred from experimental data, and beyond the technical challenges of handling massive datasets, in many cases the relevant observables are unknown (see Fig. 1).

**Figure 1.**
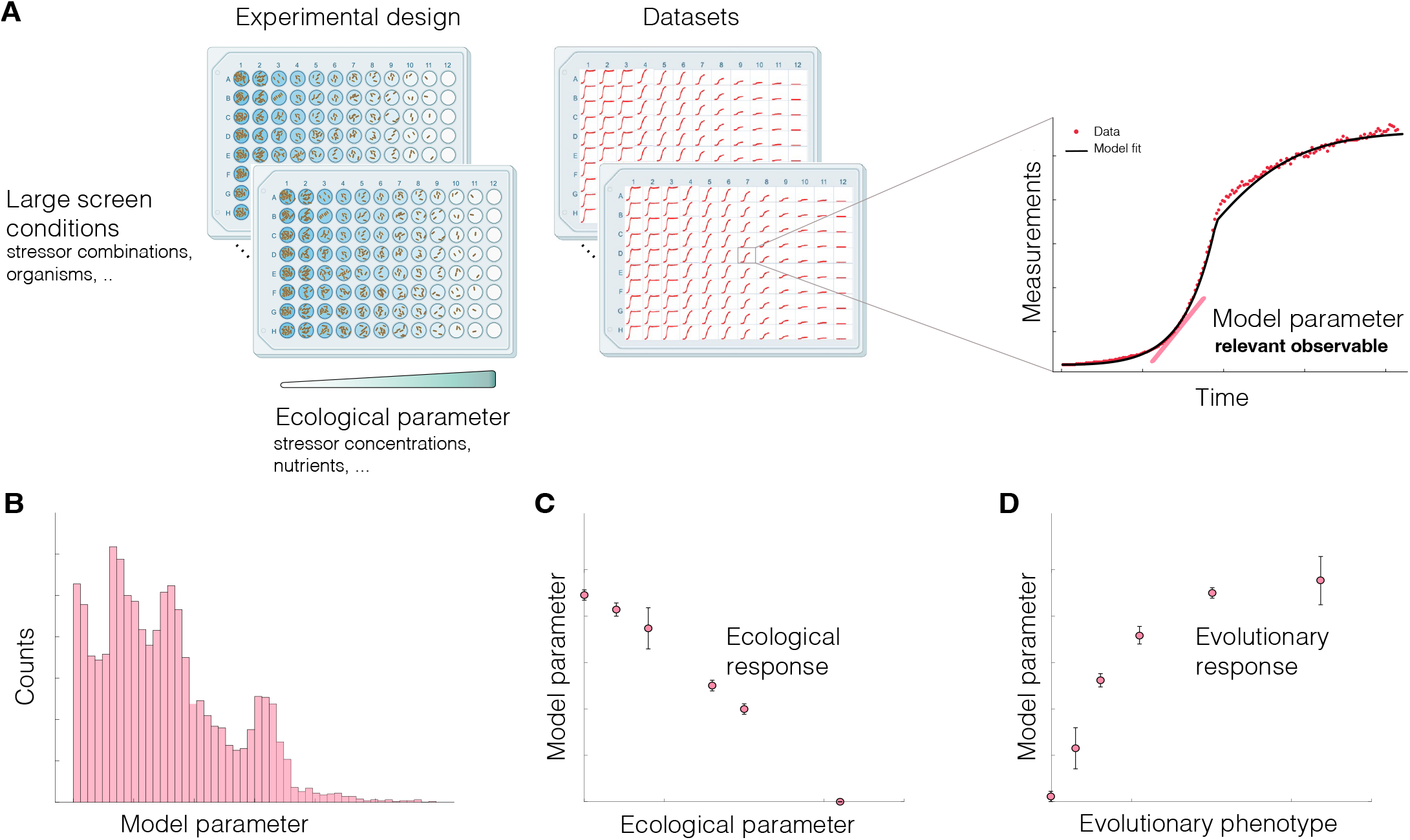
From experimental design to response patterns: **A**, Microbial kinetics experiments produce large, time-resolved datasets capturing microbial behavior across various conditions. These include ecotoxicological screens probing the effects of stressors on bacterial growth [16] or studies examining the impact of antibiotic combinations [42]. The goal is to identify relevant phenotypic markers (e.g., growth rate, total growth) by inferring these observables from microbial kinetics. This procedure enables quantitative comparison of responses across conditions. **B**, Schematic representation of a distribution of parameter values inferred from a large-scale screen: computational methods can detect regularities in these distributions, identifying similarities in the effects of perturbations within specific groups. **C**, Schematic representation of an ecological response: the relationship between model parameters and an ecological challenge defines an ecological response, such as dose-response curves for antibiotics [43] or yield-substrate relationships as a function of nutrient levels. **D**, Schematic representation of an evolutionary response: across different organisms, the relationship between inferred parameters and molecular phenotypes defines an evolutionary response. This can be used, for example, to predict antibiotic resistance evolution through metabolic fitness landscapes [44].

Available tools can assist with parameter inference for growth kinetics datasets [4–9], including more complex cases like diauxic growth [9], user-defined functions, or ordinary differential equations [5, 7], and even fitting specific dosage-response models [7, 8]. However, we lack resources to support exploratory data analysis, help identify relevant observables for further investigation and detect mathematical expressions of response patterns that can inform on biological processes underlying the observed responses. With the rapid advances in machine learning (ML), there is now a significant opportunity to enhance data analysis by combining the interpretability of model-based parameterization with the automation of ML techniques.

To establish this methodology, we introduce Kinbiont—a Julia package designed as an end-to-end pipeline for biological discovery, enabling data-driven generation of hypotheses that can be tested in targeted experiments. Leveraging Julia’s speed, flexibility, and growing popularity in biological sciences [10], Kinbiont integrates advanced solvers for ordinary differential equations (ODEs), non-linear optimization methods, signal processing, and interpretable ML algorithms. Kinbiont can fit various models of microbial dynamics, including solutions in closed-form, ODEs, and user-defined models. Unlike existing tools, Kinbiont extends model-based parameter estimation to fits with segmentation, allowing for more detailed analysis of the microbial dynamics. The inferred parameters can then be analyzed using symbolic regression [11, 12] to discover equations that capture response patterns or decision tree [13] to identify informative response patterns from large datasets.

We benchmark Kinbiont with both synthetic and real data, demonstrating its ability to fit nonstandard growth curves using publicly available datasets of diauxic curves [14] and phage-bacteria co-cultures [15]. To further illustrate Kinbiont’s potential as an integrative pipeline, we apply it to two datasets: a variation of Monod’s experiments probing auxotroph growth in response to supplemented amino acid, where symbolic regression is used to derive empirical laws; and a large-scale chemical screening dataset investigating ecotoxicological patterns [16], where decision trees explore the effects of stressors, both individually and in combination, on various phenotypic metrics of bacterial growth.

## Results

### The Kinbiont framework

Kinbiont is an open-source library freely available at https://github.com/pinheiroGroup/Kinbiont.jl or via the Julia package manager. The Kinbiont framework consists of three modules (see Fig. 2):

**Figure 2.**
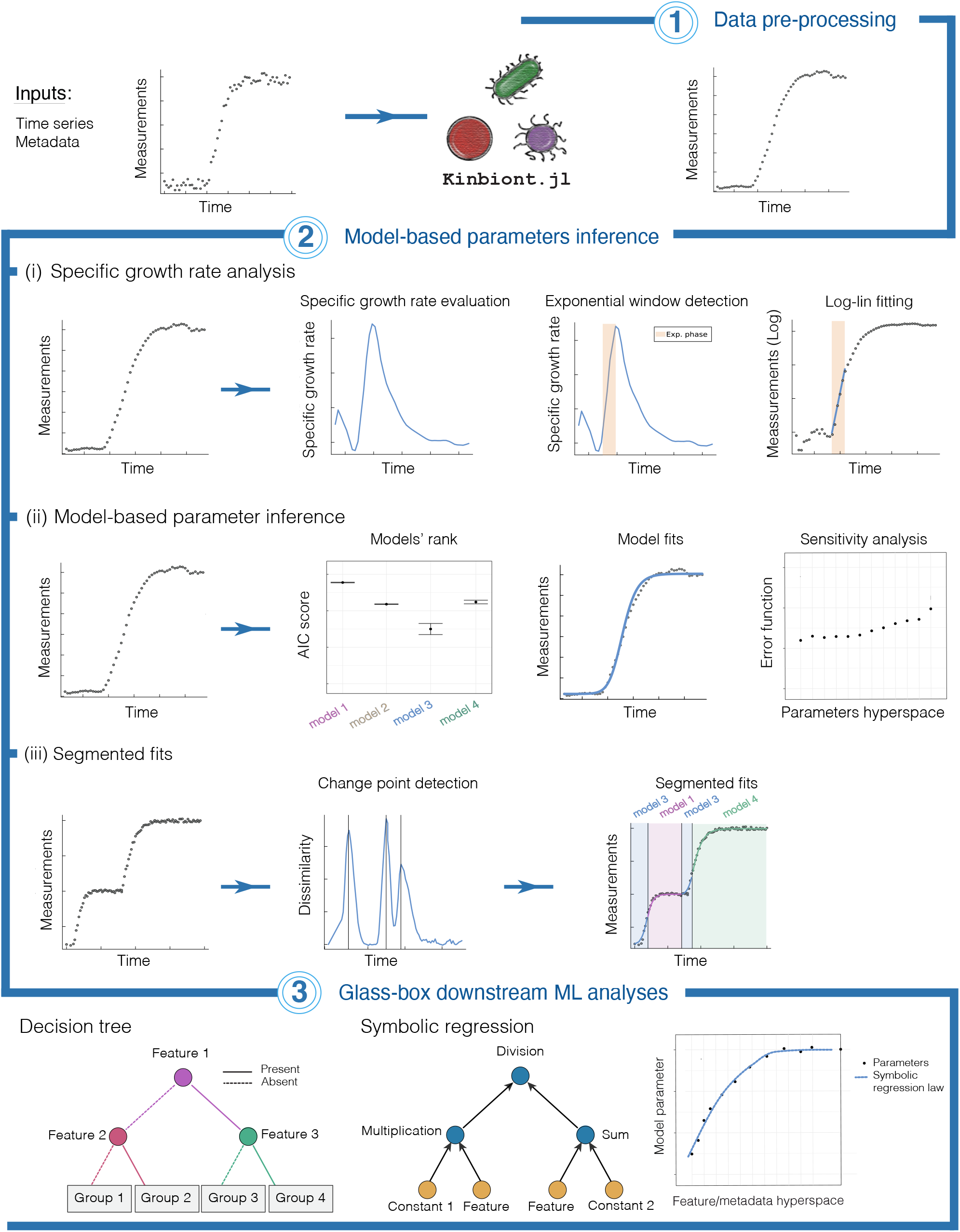
The Kinbiont framework. Kinbiont integrates three core modules: **(1) Data pre-processing**, which handles background subtraction, smoothing, and multiple scattering correction; **(2) Model-based parameter inference**, which includes methods for specific growth rate analysis, non-linear models, ordinary differential equations (ODEs), and segmented fits with change point detection; and **(3) Glass-box downstream ML analyses**, which applies interpretable ML techniques such as decision tree and symbolic regression to analyze inferred parameters for automatic detection of biological patterns.

#### 1. Data pre-processing

Kinbiont takes a time series as input with optional annotations containing features (e.g., ecological conditions or strain/species information). The input time series can undergo pre-processing based on the user’s choice, including background subtraction, averaging over replicates, multiple scattering correction for plate reader experiments [17], and data smoothing (see Methods for a detailed description of the format and pre-processing options).

#### 2. Model-based parameter inference

Kinbiont infers kinetic parameters, such as lag time, growth rate, and growth saturation, from the input time series through three distinct programs:

##### (i) Specific growth rate analysis

Focusing on exponential growth, Kinbiont identifies an exponential growth window to infer the specific or *per capita* growth rate *λ* = *d* log(*N/N*_0_)*/dt*, where *N* is the measurement and *N*_0_ is the initial value. To identify the exponential window, Kinbiont numerically generates a distribution of instantaneous specific growth rates, evaluating it locally for small intervals into which the curve is divided. Users can choose between a sliding window and a data point interpolation to define such intervals. Kinbiont uses a quantile threshold on the instantaneous specific growth rate distribution (user-defined; default set to 0.9) and returns the specific growth rate from a least squares log-linear fit in the exponential window interval (Fig. 2). Kinbiont reports the parameter of the log-lin fit with the 95% confidence interval, the exponential growth window boundaries, and the maximum exponential growth rate in the interval with the corresponding time of the maximum growth rate.

##### (ii) Nonlinear and ordinary differential equations for microbial dynamics

Kinbiont characterizes microbial dynamics by fitting a model to the time series data. Fitting is performed by solving a nonlinear optimization problem with a loss function of user choice (Methods). To handle standard and non-standard growth and death curves, we have hard-coded over 30 models, including closed-form solutions of nonlinear (NL) growth models and ODEs for microbial dynamics (Extended Data Figs. 1 - 2, Supplementary Information). Users can also define custom functions and ODE models for parameter inference.

To overcome different challenges in fitting ODE models versus analytical closed-form solutions, Kinbiont supports over 100 optimization algorithms, including global, mixed-integer, non-convex, constrained, and restart schemes [18, 19]. By default Kinbiont uses a differential evolution black box optimization with box constraints for the parameters values [18]. This option enables analysis of more general non-differentiable models and error functions, and enforces the correct range of parameter values when they are not implied in the models’ mathematical properties.

Additionally, Kinbiont provides methods for model selection [20], sensitivity analysis [21], confidence intervals estimation of inferred parameter values [22] (Methods), and it returns an output report with the optimal model parameters and the exponential growth rate. We benchmarked the accuracy of the default parameter inference using synthetic data with varying noise levels and compared the inferred parameters across different ODE and NL model representations in multiple scenarios (Methods, Figs. S2 - S3).

##### (iii) Segmented fits

To characterize non-standard growth data like diauxic curves or bacterial death and accurately capture distinct dynamical regimes, Kinbiont performs NL and ODE model fits to segmented time-series data. Kinbiont uses an offline change point detection algorithm [23] to segment the data into non-overlapping smaller time series (Methods, Supplementary Information) and initiates the fitting procedure. This program performs model selection for each segment while ensuring the continuity of the solution. Kinbiont also offers flexibility in selecting the number of change points by automatically ranking models with different segment numbers within a user-defined range.

Along with fitting routines, Kinbiont includes simulators of microbial dynamics to generate synthetic datasets with either ODEs or a stochastic Poisson process (Methods).

#### 3. Glass-box downstream machine learning analysis

To aid hypothesis generation and biological discovery, Kinbiont integrates interpretable ML methods such as symbolic regression and decision tree. Applied to model parameters extracted from large datasets, these methods enable the detection of mathematical expressions of response patterns, ranking the importance of features like stressors, and provide graphical knowledge representations.

### Kinbiont enables model-based parameter inference in non-standard microbial growth kinetics

To showcase Kinbiont’s performance in analyzing non-standard microbial growth kinetics, we applied it to a dataset addressing diauxic shifts in multi-resource environments [14]. Diauxie occurs when microbes sequentially utilize different nutrient sources. Starting with a typical single-resource growth curve, diauxic growth features a second lag phase that reflects the time needed for metabolic rearrangements to process the second nutrient. This phase is then followed by a new period of exponential growth until the culture reaches saturation [14]. The duration of the lag phase, the exponential growth rates, and the saturation values are relevant for quantitatively understanding microbial responses in multi-nutrient environments. However, we lack automatic tools for extracting these parameters using model representations, and current methods often require manual curation [7].

We applied segmented fits to the optical density (OD) time series of *Acinetobacter* species growth under various nutrient combinations, using the ODEs from the exponential (Eq. 1), logistic (Eq. 2), heterogeneous population (HPM; Eq.3), and exponential HPM (Eq. 4) models (see Methods for the explicit expressions of these equations and details of the fitting procedure). As illustrated in Fig. 3 **A, B**, Kinbiont accurately fits the entire dataset, reproducing the measured *Acinetobacter* growth curves from Ref. [14] across the entire range of conditions.

**Figure 3.**
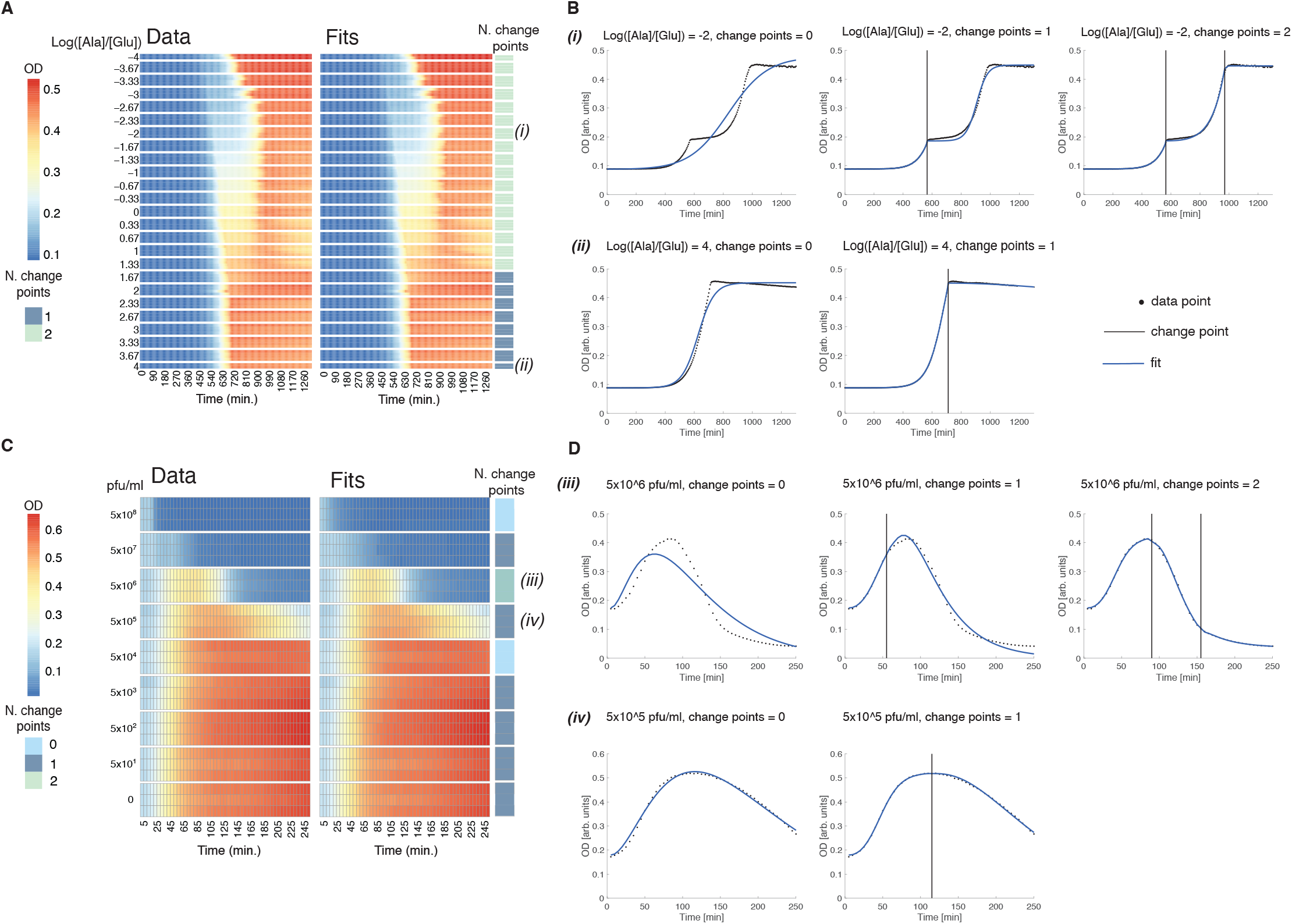
Fitting non-standard kinetics: **A**, Heatmaps showing time-resolved optical densities (OD) for diauxic growth experiments from Ref. [14] (left), paired with Kinbiont fits for each condition of [Ala*/*Glu] concentration ratios, determined through a direct search to find the optimal number of change points (right). The optimal number of change points is indicated for each case (Fig. S5 shows the AICc estimator for fits with different numbers of change points). **B**, Sample fits displaying the best fits with an increasing number of segments, up to the optimal fit; vertical lines indicate the change point locations. **C**, Heatmaps showing time-resolved optical densities (OD) for the dataset from Ref. [15], which explores the interaction between bacteria and phages at varying phage inoculum concentrations (pfu/ml) (left), paired with Kinbiont fits for each condition determined through a direct search to find the optimal number of change points (right) (Fig. S6 shows the AICc estimator for fits with different numbers of change points). Sample fits are shown in **D**, with different numbers of change points up to the optimal segmented fit.

Next, we applied Kinbiont to optical density (OD) data from *Escherichia coli* cultures infected with various concentrations of T4 phage [15]. In viral infections of bacterial cultures, the growth kinetics exhibit distinct phases of growth and death; the latter resulting from cell lysis, which is closely related to the bacteriophages inoculum concentration. As a result, these time series are qualitatively different from those observed in standard growth experiments or the diauxic growth discussed above.

For this analysis, we considered the ODEs of the following models: exponential (Eq. 1), logistic (Eq. 2) HPM (Eq. 3), the exponential HPM (Eq. 4), and HPM with inhibition and death (Eq. 5) (see Methods for the explicit expressions of these equations and details of the fitting procedure). As shown in Fig. 3 **C, D**, Kinbiont is able to accurately fit the growth curves of *E. coli* across different phage exposure conditions, successfully reconstructing the measured data of Ref. [15].

Together, these case studies demonstrate Kinbiont’s applicability beyond standard growth kinetics, showcasing its ability to automatically reconstruct piece-wise models for a variety of microbial kinetics profiles. Ecological response models constructed with parameters inferred from these growth curves can address fundamental questions in microbiology, including microbial coexistence in multi-resource environments (cf. analysis in Ref. [14]) and the effects of phage-bacteria interactions on bacterial growth inhibition (cf. analysis in Ref. [15]).

### Kinbiont identifies mathematical models underlying microbial responses

To evaluate Kinbiont as an integrative pipeline, we performed a variation of Monod’s classical experiment on limiting nutrients [1]. Using an *E. coli* auxotroph that cannot synthesize the amino acid methionine (Δ*metA* knockout), we recorded growth kinetics in minimal media with different concentrations of supplemented methionine using OD measurements in batch culture (Methods).

The data were preprocessed with background subtraction and multiple-scattering correction (Methods). Although sometimes overlooked, this step is essential: neglecting background effects can lead to misestimation of the exponential growth rate (Extended Data Fig. 3 **A**), while ignoring multiple scattering events can result in underestimated saturation levels (Extended Data Fig. 3 **B**). We applied a segmented fitting approach with one change point, considering the logistic (Eq. 2) and exponential HPM (Eq. 3) models for every ecological condition with different concentrations of supplemented methionine (Fig. S7). We extracted the exponential growth rate of the first segment and used the saturation OD of the second segment to account for the total growth. The response profiles of both observables as a function of methionine concentration are shown in Fig. 4.

**Figure 4.**
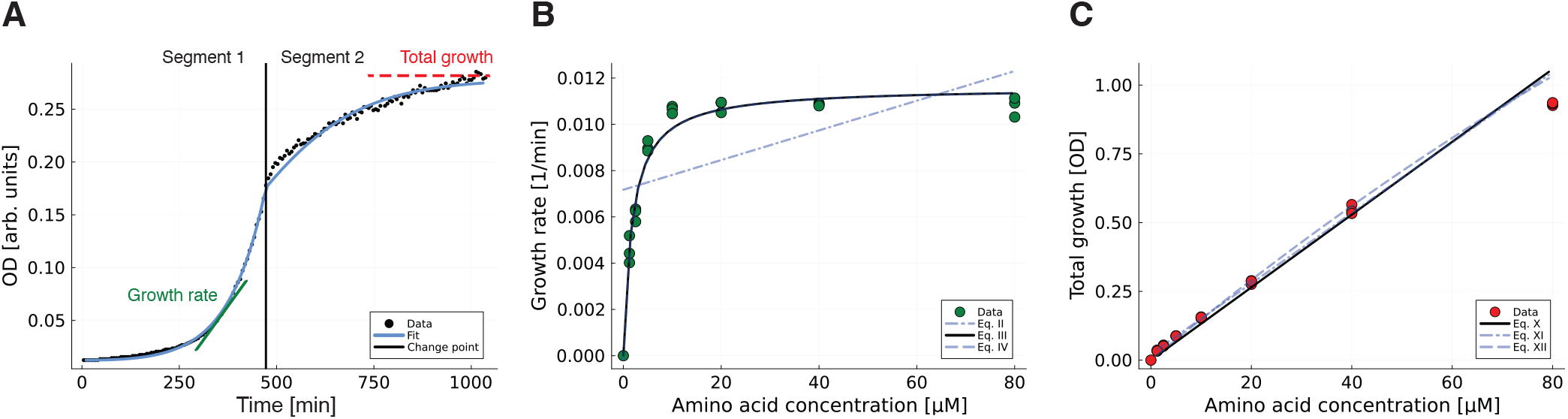
Automated detection of empirical responses: **A**, Sample growth curve illustrating the fitting procedure and parameter inference from the optical density (OD) time series of the methionine auxotroph strain. This example corresponds to 10*µ*M of supplied methionine; all growth curves and corresponding optimal fits are shown in Fig. S7. The exponential growth rate and total growth were extracted from the first and second segments, respectively. **B**, Inferred growth rates at different concentrations of methionine (dots), together with plots of a subset of the equations from the hall-of-fame of symbolic regression. The solid black line represents the Monod equation. **C**, Inferred total growth at different methionine concentrations, accompanied by plots with candidate equations from the hall-of-fame of symbolic regression. The substrate-yield relationship for the total growth response is again shown as the solid black line. The expressions for all equations in the hall-of-fame of symbolic regression are provided in Extended Data Fig. 4.

Next, we employed symbolic regression [11] to find mathematical expressions matching the responses of growth rate and total growth with the change in amino acid concentration (Methods). The proposed laws, their complexity scores, and mean squared errors are presented in Extended Data Fig. 4. Remarkably, the algorithm successfully identified the celebrated empirical relationships discovered by Monod [1] within the hall-of-fame of candidate models for both observables. As shown in Fig. S8, these results were replicated for another methionine auxotroph strain.

This result establishes a proof-of-concept for empirical law discovery related to ecological perturbations such as changes in nutrient quality or concentration.

### Kinbiont ranks the impact of stressors on growth-phase-specific parameters

A different class of problems in ecology concerns feature identification. Microbes are exposed to multiple environmental stressors, such as antibiotics or pollutants. But how do interactions among stressors impact microbial physiology and ecosystem ecology? [24]

To address this question, we applied Kinbiont to the dataset from Ref. [16], which recently profiled the growth kinetics of 12 bacterial isolates: two strains of *P. baetica*, two strains of *A. humicola*, one strain of *F. glacei, R. herbae, S. faeni, C. gallinarum, A. popoffii, N. soli, E. coli*, and *A. fisheri*, and a mixture of 10 of them. These isolates were tested under every possible combination of eight different pollutants known to be prevalent in freshwater environments: Amoxicillin, Tebuconazole, Metaldehyde, Oxytetracycline, Chlorothalonil, Imidacloprid, Diflufenican, and Glyphosate (255 chemical stressor mixtures in total, amounting to 15120 growth curves in addition to controls). After background subtraction, we fit the entire dataset with the Richards model (Eq. 6 in Methods) to extract the duration of the lag phase, the exponential growth rate, and total growth (about 10% of the curves with an average relative error larger than a threshold were discarded from further analyses; Methods, Fig. S9).

For each of these observables, we performed a decision tree regression with splitting criteria based on the Gini impurity. We evaluated the model’s performance by considering different tree depths and using a 10-fold cross-validation for each case (Methods, Figs. S10 - S12). Interestingly, the coefficients of determination indicate that the effect of a given chemical is not necessarily relevant across all observables of a given strain. For example, we obtained a cross-validation coefficient of determination *R*^2^ *>* 0.5 for all observables of *N. soli*, but only for the growth rate of *A. fischeri* (Methods, Figs. S10 - S12).

Next, we ranked stressors for each strain and model parameter using the impurity importance score, based on the largest shifts in mean parameter distributions in response to the presence or absence of each stressor. The score values (Fig. 5) indicate that the response to a chemical is highly species-specific. More surprisingly, they reveal an intra-strain differential response across the various phases of growth (Fig. 5 **A, C, E**). For example, Imidacloprid significantly affects the growth rate in the species *N. soli*, while the total growth is mostly affected by Oxytetracycline, and the duration of the lag phase is mainly affected by Tebuconazole. Importantly, summary metrics like the commonly used area under the curve (AUC) cannot resolve growth-phase specific stressor effects. As shown in the Extended Data Fig. 5, the AUC primarily serves as a proxy for the total growth of cultures over a fixed time interval. However, a given AUC value can be highly degenerate with respect to growth rate or lag phase duration (Extended Data Fig. 5 **A, C**).

**Figure 5.**
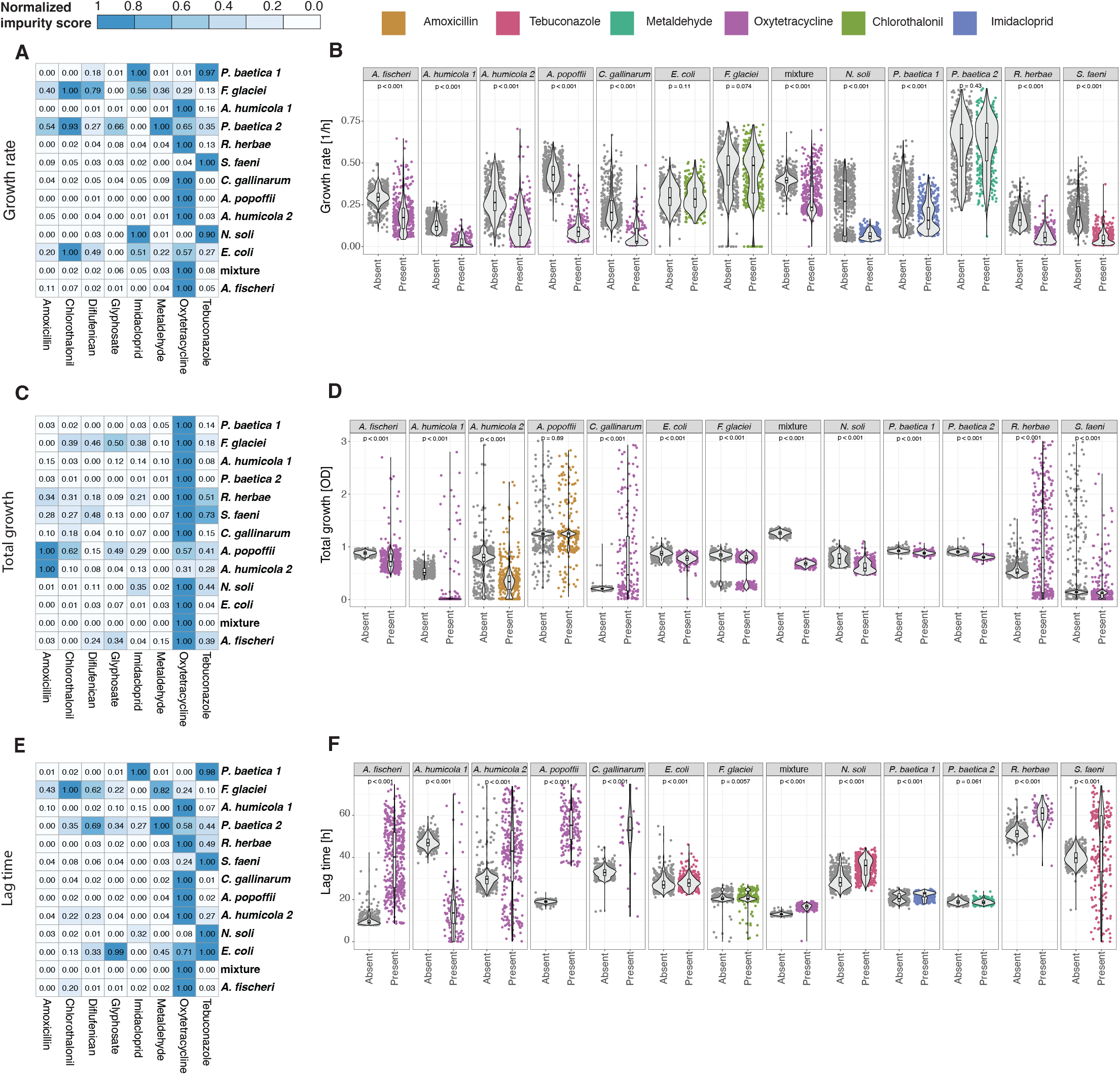
Characterization of chemical perturbations: **A, C**, and **E**, Heatmaps displaying the normalized impurity scores (scaled between 0 and 1) for each chemical, strain, and kinetic parameter from the decision trees evaluated at maximum depth (Methods). High values indicate that the growth property is perturbed by the chemical, while low values suggest that the strain’s kinetic parameter remains unaffected. **B, D**, and **F**, Distribution splitting analysis for each strain and parameter, illustrating the effect of the presence or absence of the most relevant chemical (indicated by the highest impurity score). Each panel includes the p-value from the Wilcoxon test, indicating the statistical significance of the observed differences.

More broadly, the methodology presented here can be applied to the design of targeted interventions, aimed at achieving specific effects on parameters of microbial growth kinetics like biomass production optimization and therapy selection.

### Kinbiont resolves interactions in combinations of stressors

Beyond ranking the relevance of stressors in chemical perturbation screens, the graphical models in Kinbiont can characterize the impact of stressor combinations. To illustrate this analysis, we consider the decision trees of *N. soli* and the mixture of strains, which display a consistent coefficient of determination *R*^2^ ≳ 0.5 across all the different observables (Methods, Figs. S10 - S12). Figure 6 shows the first two levels of the maximum depth trees for each kinetic parameter, together with the distribution of parameter values at each leaf (Methods). Notably, all chemicals displayed at this level significantly split the distributions (p-values from Wilcoxon test), confirming the decision tree’s ability to resolve the chemical landscape beyond the single stressor with highest effect.

**Figure 6.**
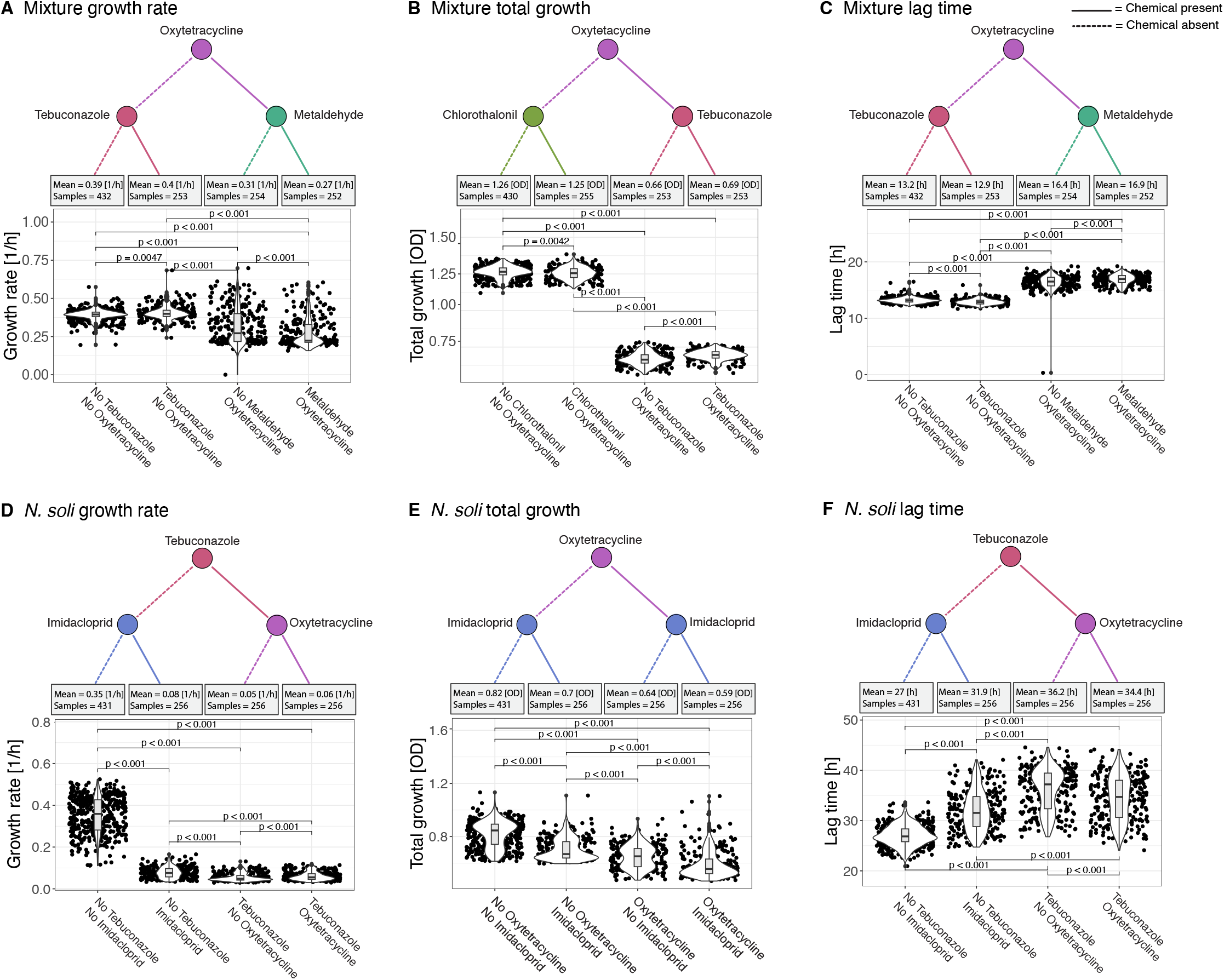
Profiling stressor combinations with Kinbiont: Graphical representation of the first two levels of the maximum depth decision trees for the different kinetic parameters inferred for the mixture of strains **(A - C)** and for the strain N. soli **(D - E)**. The dashed lines indicate that the chemicals present in the parent nodes are absent in the next level of the tree, while the solid lines indicate that the chemicals are present. Violin plots at each leaf show the corresponding distributions, with p-values from the Wilcoxon test between each pair of distributions quantifying the significance of the splitting with respect to the distribution mean. In addition to identifying the chemicals with the most significant statistical effects - like the pronounced impact of Oxytetracycline on the total growth of the mixture in panel **B** — this graphical representation can also be used to explore chemical interactions. For instance, it can detect synergistic interactions between Oxytetracycline and Imidacloprid in **E**, and antagonistic interactions between Tebuconazole and Oxytetracycline in **D** and **F**.

Expanding on the graphical representation of decision trees, Kinbiont can be used to identify synergistic or antagonistic interactions. Starting with the isolates mixture (Fig. 6 **A** - **C**), we observe a weak antagonistic effect between Oxytetracycline and Metaldehyde in the distribution of growth rates, leading to a mean increase in generation time higher than 10% (mean doubling time gain of 20 mins; Fig. 6 **A**). In *N. soli*, the distribution of growth rates at the leaf with Tebuconazole but no Oxytetracycline is shifted to lower values compared to the leaf where both chemicals are present, indicating a small antagonistic effect between these drugs on the growth rate (Fig. 6 **D**). While this combination of chemicals exhibits a similar antagonistic effect on the lag phase of this strain (Fig. 6 **F**), the total growth of *N. soli* is synergistically affected by the presence of Oxytetracycline and Imidacloprid, as the presence of both drugs results in a distribution with a lower mean (Fig. 6 **E**).

These results illustrate how Kinbiont can be used to generate testable hypotheses about chemical interactions from large datasets to inform combined interventions. For example, we can use it to identify the statistically most relevant interactions for further experimentation, use the findings to interrogate unknown modes of action of stressors based on systematic comparisons with compounds that have already been characterized, and reduce the search space for possible stressor targets.

## Discussion

Kinbiont is an open-source library designed to accelerate biological discovery by enhancing microbial time-series data analysis through an integrated approach that combines the interpretability of microbial dynamics models with the automation of machine learning techniques. It processes microbial time-series data and generates testable hypotheses on ecological or evolutionary responses. To achieve this, Kinbiont integrates independent modules for data pre-processing, model-based parameter inference, and downstream analysis using machine learning algorithms. To our knowledge, Kinbiont is the first tool to integrate downstream analyses with machine learning methods in microbial kinetics.

Kinbiont introduces various innovations that expand its utility beyond conventional methods for parameter inference. First, it leverages Julia’s extensive ecosystem of numerical methods, providing a significant performance advantage for computationally intensive tasks [10]. Second, Kinbiont offers the most comprehensive collection of pre-built microbial dynamics models and supports custom models in both nonlinear (NL) and ordinary differential equation (ODE) representations. Third, a key innovation is the introduction of segmented fits – a method that combines signal analysis, model selection, and optimization-based fitting for time-series data with multiple growth phases. As demonstrated by analyzing diauxic shifts and phage infections, segmented fits allow for accurate extraction of growth-phase-specific parameters in non-standard microbial growth patterns.

In addition, Kinbiont integrates machine learning into the pipeline through methods like symbolic regression and decision tree. Symbolic regression automates the discovery of mathematical relationships between microbial observables, extending the pioneering work of Monod to modern datasets. This enables the identification of empirical laws and patterns in microbial responses across diverse conditions. By combining experiments and data analysis with the Kinbiont framework, we established the first proof of principle of automatic law detection in microbiology.

Decision tree is a complementary ML method implemented to identify key features that affect microbial behavior. As demonstrated in our analysis of the ecotoxicological response study [16], the combination of model-based parameter inference and decision trees can detect species-specific and growth-phase-specific effects of stressors. These insights could guide targeted treatments, such as identifying chemicals or their combinations that selectively impact overall bacterial growth or growth rate. While we did not apply Kinbiont to evolutionary response data due to a lack of available datasets, similar analysis can be performed for evolutionary studies by using different genotypes as features or modeling changes in the variants’ phenotypes.

Looking ahead, the high modularity of Kinbiont will support the integration of additional methods for downstream analysis, such as explainable boosting machines [25] and non-linear mixed-effects models [26]. These methods are well-suited for handling high-dimensional data and can be tailored to explore pharmacological studies and interactions between features. This will further expand Kinbiont’s utility in fields such as synthetic biology, where it can optimize gene circuits by identifying genetic modifications that enhance system stability; metabolic engineering, where it can analyze how perturbations of metabolic pathways influence product yield; and pathogen-host interactions, where it can help identify how specific host genes affect infection outcomes.

Kinbiont establishes a new methodology in microbiology by not only assisting with data analysis but also aiding in theory formulation. Its modular, extensible design allows it to adapt to evolving computational techniques and emerging research needs, making it a long-term tool for scientific discovery. This flexibility positions Kinbiont as a powerful resource for modern microbiology research.

## Methods

### Input Data

Kinbiont can be used within a Julia notebook or to work directly with data files. In a Julia notebook, Kinbiont expects input data as a 2*×n* matrix, where *n* is the number of time points. The first row accounts for the time values, and the second row contains the measured quantity, such as optical density (OD) or colony-forming units.

Alternatively, Kinbiont offers APIs for handling external .csv files. Users can specify paths to a primary data file and, optionally, an annotation file. For datasets containing multiple time series, the data file should be organized as a matrix. The first row must include the time series’ identifier, followed by rows of numerical measurements across time points. The first column is reserved for the time values, while the subsequent columns keep the data for each individual case. If an annotation file is provided, it should be a two-column .csv: the first column lists the names of the time series, and the second column includes a unique identifier for biological replicates, with an optional exclusion flag.

Detailed data format descriptions and examples are available in the GitHub repository [27].

### Pre-processing of bacterial growth curves

Kinbiont provides the following pre-processing options to prepare data before model fitting:

#### 1. Background subtraction, correction for negative values, and averaging of replicates

If an annotation file is provided, Kinbiont can automatically subtract blank values by calculating a mean for all blanks or by generating a rolling time average to subtract from the data longitudinally. Negative values resulting from this subtraction, which may complicate fitting procedures such as log-lin fitting, are handled by either removing them or imputing alternative values from a user-defined threshold or the distribution of blank values. Additionally, Kinbiont can average data from wells with the same unique identifier if annotations are provided.

#### 2. Data smoothing

Before inference, users can smooth the data using either a rolling average with a user-defined window size or locally weighted scatterplot smoothing (LOWESS) [28].

#### 3. Correction for multiple scattering in OD measurements

Analyses of OD measurements often assume that OD values are directly proportional to cell count, but this assumption holds only under specific conditions [17]. To correct for deviations due to multiple scattering, Kinbiont allows users to input a calibration curve. This consists of a two-dimensional matrix where the first dimension contains raw OD readings and the second dimension contains OD values obtained from the same samples with multiple scattering free measurements to be used as a reference [29] (cf. section Multiple scattering calibration data).

Kinbiont offers two methods for correcting the growth curve datapoints:

- Interpolation: using the Steffen monotonic interpolation algorithm [30].
- Extrapolation: by fitting the following exponential model from [29]:

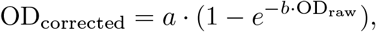

where *a* and *b* are fit parameters.

When using interpolation, it is important that the calibration data covers the entire growth curve range to avoid incorrect imputation of values outside this range.

### List of models used in this study

The models listed below were used in the analyses presented in this study and are referenced throughout the text. A complete list of models implemented in Kinbiont is provided in Extended Data Figs. 1 and 2, with detailed parameter descriptions available in the Supplementary Information. Implementation details can also be found in the GitHub repository [27].

#### ODE models

- Exponential:

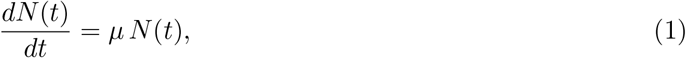

where *µ* is the growth rate.
- Logistic [31]:

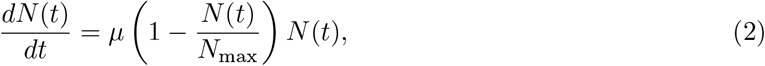

where *µ* is the growth rate, and *N*_max_ the total growth.
- Heterogeneous Population Model (HPM) [32]:

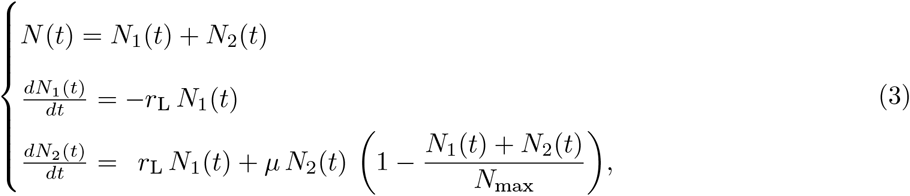

where *N*_1_ is the population of dormant cells, *N*_2_ is the population of active cells capable of duplicating, *µ* is the growth rate, *N*_max_ is the total growth, and *r*_L_ is the lag rate, defined as the rate of transition between the *N*_1_ and *N*_2_ populations. Here, we assume that all cells are in the dormant state at the start (i.e., *N*_1_(*t* = 0) = OD(*t* = 0), and *N*_2_(*t* = 0) = 0).
- Exponential HPM:

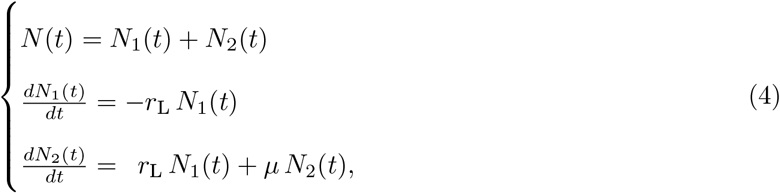

where similarly to the HPM model, *N*_1_ and *N*_2_ refer to the populations of dormant and active cells, respectively. *µ* is the growth rate, and the lag rate *r*_L_ denotes the transition between the *N*_1_ and *N*_2_ populations. Here, we also assume that all cells are in the dormant state at the start (i.e., *N*_1_(*t* = 0) = OD(*t* = 0), and *N*_2_(*t* = 0) = 0).
- Heterogeneous Population Model with Inhibition and Death :

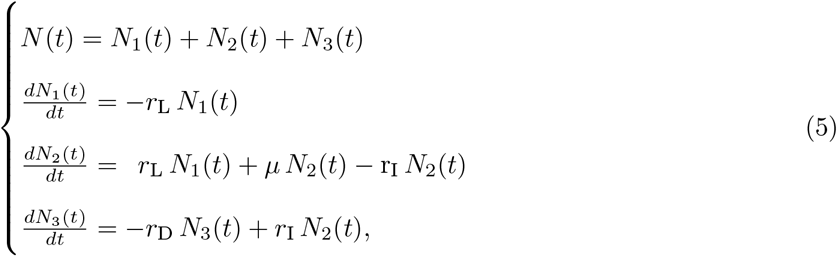

where *N*_1_, *N*_2_, and *N*_3_ refer to the populations of dormant, active, and inhibited cells. In contrast to the population of actively duplicating cells in *N*_2_, cells in the *N*_3_ inhibited state are active but do not duplicate. The other parameters are the growth rate *µ*, the lag rate for the transition between *N*_1_ and *N*_2_, *r*_L_, the lag rate for the transition between *N*_2_ and *N*_3_, *r*_I_, and the rate at which cells die, *r*_D_. As in the cases above, we assume that all cells start in the dormant state (i.e., *N*_1_(*t* = 0) = OD(*t* = 0), *N*_2_(*t* = 0) = *N*_3_(*t* = 0) = 0).

To fix the initial condition when fitting differential equations, use *N* (*t* = 0) = OD(*t* = 0).

#### NL models

- Richards model [33]:

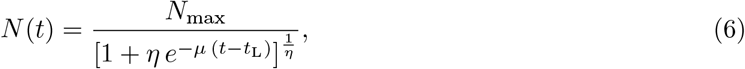

where *µ* is the growth rate, *N*_max_ is the total growth, *t*_L_ the lag time, and *η* a shape constant.

### Parameters inference as a nonlinear optimization problem

In Kinbiont, parameter inference is framed as a nonlinear optimization problem. Given a set of parameters *P*, a time series of data points *N* (*t*_*i*_) where *i* = 1, …, *n*, and 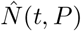, representing the proposed bacterial kinetics (e.g., an approximate numerical solution of one of the ODEs evaluated at time *t* with parameters *P*), we define an objective function:

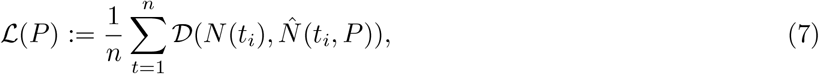

where 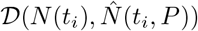 represents the distance between the data and the numerical solution. This formulation transforms parameter inference into an optimization problem aimed at finding a set of hyperparameters *P* ^*∗*^ that minimizes the objective function:

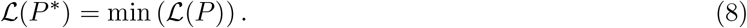

Kinbiont uses the package `Optimization.jl` [18] which integrates over 100 optimization schemes, including local and global optimization methods, derivative-free methods, gradient-based approaches, constrained optimization, and multi-start techniques. This variety allows users to select the most appropriate method based on their time constraints, the complexity of the loss function, or specific performance requirements. Different loss functions are hard-coded in Kinbiont, such as the *L*2 norm and the relative error. In the analyses of the main text, we used the latter, defined as

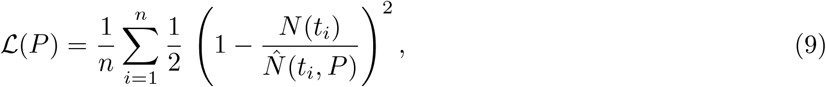

where *n* is the number of data points. A comprehensive list of available loss functions can be found in the Supplementary Information and the GitHub repository [27].

### Model selection

Selecting an appropriate growth model for bacterial kinetics can involve trial and error due to the non-mechanistic nature of macroscopic models [34]. To streamline this process, Kinbiont features a model selection tool based on the Akaike Information Criterion (AIC) [20].

Kinbiont fits all models from a user-defined list and evaluates the AIC using the formula [20]:

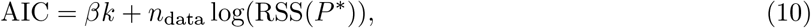

where *k* is the number of model parameters, *β* is a user-defined penalty, *n* is the number of data points, and RSS is the sum of root squared errors for the fits with optimal parameters *P* ^*∗*^. Kinbiont returns the AIC values for all models and identifies the model with the lowest AIC as the “best” fit. Users can also opt to calculate a corrected AIC for small sample sizes using:

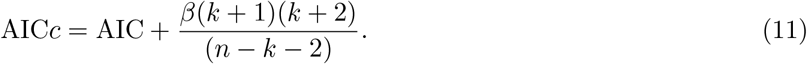

### Change point detection algorithm

Kinbiont implements two offline change point detection methods [23]: (*i*) a sliding window with linear fitting that minimizes the absolute deviation from the mean of the data, and (*ii*) the least square density difference (LSDD) method [35]. The key distinction between these methods lies in the cost function **C**, used to evaluate the dissimilarity curve **D**,

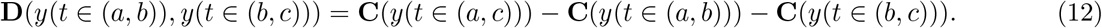

In method (*i*), the cost function is computed as:

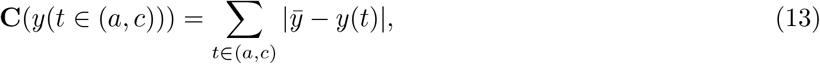

where *t*_0_ *≤ a < c < b ≤ t*_end_, *y*(*t*) is the theoretical prediction of a linear fit in the considered segment, and 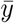 is the average signal in the considered time interval. Method (*ii*), uses the Julia package `ChangePointDetection.jl` to evaluate the *L*2 norm for the difference between the probability densities estimated on the datapoints of two adjacent segments [35]. The change point detection process involves:

#### 1. Dissimilarity curve construction

The algorithm constructs a dissimilarity curve using a sliding window, comparing adjacent windows along the time series (or its derivative, if specified by the user).

#### 2. Peak detection

A peak detection algorithm identifies significant changes in the dissimilarity curve, corresponding to potential change points in the growth dynamics. The user specifies the desired number of change points (*n*_CP_), and the algorithm returns the peak positions ranked by prominence.

### Fitting growth models with segmentation

After detecting the change points, Kinbiont fits the data using one of two approaches, depending on whether the user selects ODE or NL models. For ODE models, Kinbiont employs a forward-fitting approach. The process begins with the first segment, where the model selection algorithm identifies the “best” ODE model from a user-defined list, based on the AIC criterion. Subsequent segments are then fitted, ensuring continuity by enforcing the initial condition at the start of each segment. For NL models, the fitting procedure is performed in reverse order. Continuity across segments is maintained by incorporating a penalty term in the loss function to ensure consistency between segments.

### Selection of the number of change points

When the exact number of change points (*n*_CP_) is unknown, Kinbiont provides an automatic selection feature. Kinbiont uses a direct search algorithm to evaluate all possible segmented models, given a maximum number of change points 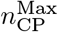. The algorithm generates all combinations of change points, ranging from 0 to 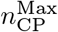, fits each combination, and selects the model with the minimal AIC (or AICc). The selection process incorporates a linear penalty on the number of segments:

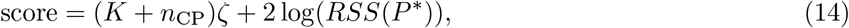

where *K* is the total number of parameters across all differential equations used, *n*_CP_ is the number of change points, and *ζ* is a smoothing parameter as described in [23].

The choice of *ζ* influences model selection: lower values of *ζ* favor models with more change points, potentially capturing complex dynamics, while higher values favor simpler models with fewer change points, emphasizing parsimony.

This approach enables Kinbiont to automatically determine the optimal number of change points, ensuring data-driven and accurate segmentation of the time series. The flexibility in setting *ζ* allows users to balance model complexity with fit quality, adapting the analysis to the specifics of the data.

### Sensitivity analysis

Kinbiont includes an automatic sensitivity analysis feature to ensure that the optimization algorithm produces stable inferences, regardless of parameter initialization. This is accomplished using the Morris method for global sensitivity analysis [21]. The Morris method offers a simple and computationally efficient way to identify important input variables, especially in models with a large number of parameters. During each iteration, the method adjusts one input parameter at a time — specifically, one of the initial guesses — and then reevaluates the algorithm’s output. This approach highlights the input variables that have the most significant influence on the model, indicating which ones may require more detailed analysis.

### Confidence interval estimation for non-linear fits

When fitting data with a non-linear function, confidence intervals for the inferred parameters, as well as their means, can be estimated using bootstrap or Monte Carlo methods. In the bootstrap method, the user specifies the number of fit repetitions and the portion of data for each fit. The data are randomly sampled and fitted multiple times. The 95% confidence interval and mean of each parameter are then calculated from the distribution of the objective functions [22]. In the Monte Carlo approach, the user specifies the number of fit repetitions, and for each fit, random noise from the empirical noise distribution of the blanks (with zero mean) is added to the data. The 95% confidence intervals and parameter means are then derived from the distribution of fit results [36].

### Interpretable machine learning methods for downstream analyses

Kinbiont incorporates different interpretable machine learning methods. An overview of these methods is given below, with specific application details provided in the relevant sections of this paper.

#### Decision Tree

`DecisionTree. jl` is a Julia implementation of the classification and regression tree algorithm as described in Ref. [13]. This method constructs a graphical model, commonly referred to as a decision tree, by recursively partitioning the dataset and applying a simple prediction model within each partition. For classification trees, which handle categorical variables, prediction accuracy is based on misclassification costs. Regression trees, used for continuous or ordered discrete variables, measure accuracy by the squared difference between observed and predicted values. This method has been applied in previous studies evaluating the impact of media chemical composition on bacterial growth [37].

#### Symbolic Regression

`SymbolicRegression. jl` is a Julia implementation of symbolic regression as described in Ref. [11]. Symbolic regression aims to produce an interpretable model represented as an analytical function. In this approach, equations are organized as trees, with leaves corresponding to constants or dataset features, and internal nodes representing operations. Binary operations, like scalar products, have two daughter nodes, while unary operations, such as logarithms, have one. The algorithm searches for an expression that minimizes mean square error while maintaining simplicity, penalizing complexity based on the number of nodes in the tree. This search is conducted using a multi-population evolutionary algorithm, where different populations (ensembles of trees) evolve asynchronously [38].

### Parameter inference in non-standard microbial kinetics

The *Acinetobacter* dataset from Ref. [14] consists of 96 background-corrected curves growth profiling 25 different combinations of Alanine and Glutamate concentrations, with a logarithmic ratio of concentrations 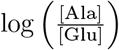 spanning the range from -4 to 4, sampled in increments of 0.333. Most conditions have four replicates, except for 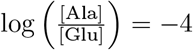 and 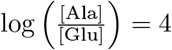, which have two replicates each.

In preparation for the change point detection, we applied a smoothing filter to the growth curves using a rolling average over three time points and then employed a linear change point detection algorithm on the specific growth rate, which was evaluated using a sliding window of six points. The change point detection was performed with a window size of ten points, and the optimal number of change points was determined through a direct search within the range of 0 to 5. For each segmented time series, we fitted the following ODE models by minimizing the relative error loss function (Eq. 9): the exponential model (Eq. 1), the logistic model (Eq. 2), the HPM (Eq. 3), and the exponential HPM (Eq. 4). Model selection was carried out using the corrected AIC score (Eq. 11), with a penalty parameter set to 2. Corrected AIC scores for all model fits are shown in Fig. S5.

The phage-bacteria interaction dataset from Ref. [15] consists of growth curves for *E. coli* initially inoculated at 10^8^ cfu/ml, with various concentrations of T4 phages added to the wells. Each concentration was tested in triplicate, resulting in 28 growth curves: 5 *×* 10^8^ pfu/ml, 5 *×* 10^7^ pfu/ml, 5 *×* 10^6^ pfu/ml, 5 *×* 10^5^ pfu/ml, 5 *×* 10^4^ pfu/ml, 5 *×* 10^3^ pfu/ml, 5 *×* 10^2^ pfu/ml, 5 *×* 10 pfu/ml, and 0 pfu/ml. We pre-processed the data by subtracting the blanks and smoothing the curves using LOWESS (with default parameters). Subsequently, we applied a linear change point detection algorithm to the specific growth rate, which was evaluated using a sliding window of six points, with an additional window size of eight points for the change point detection. A direct search was conducted to identify the optimal number of change points, ranging from 0 to 3. For each segment, we fitted ODE models by minimizing the relative error loss function (Eq. 9): the exponential model (Eq. 1), the logistic model (Eq. 2), the HPM (Eq. 3), the exponential HPM (Eq. 4), and the HPM with death model (Eq. 5). Model selection was carried out using the corrected AIC score (Eq. 11), with a penalty parameter set to 2. Corrected AIC scores for all model fits are shown in Fig. S6.

### Monod experiment data generation

M9 Medium was prepared by diluting a 5X stock solution of M9 salts (Sigma Aldrich) 1:5 in deionized (milliQ) water. The medium was then supplemented with 0.24 g/L MgSO_4_, 0.011 g/L CaCl_2_ and 0.36 g/L D-glucose as the sole carbon source. L-methionine was added to reach concentrations between 3.125 and 80 *µ*M. *E. coli* K-12 MG1655 Δ*metA* knockouts were used as methionine auxotrophs (the strains were a gift from Martin Ackermann’s Lab, ETH). Each strain was stored as *−*80°C glycerol stock, streaked on LB-Agar plates, and grown overnight at 37°C. The following day, a single colony was picked and inoculated in 15 mL of LB broth in a 100 mL Erlenmeyer flask. The liquid culture was grown overnight at 37°C with 150 rpm orbital shaking. Once saturation was reached, OD was measured with a ThermoFisher Genesys spectrophotometer, and the culture was diluted to an initial OD of 0.01 for microplate experiments.

Growth experiments were performed at 37°C with 800 rpm orbital shaking in 48-well microplates, each filled with 1 mL of culture, and OD (600 nm) measurements were taken every 6 minutes using a BioTek Synergy H1 microplate reader. Each experiment included at least two blank wells containing only growth media.

### Multiple scattering calibration data generation

This calibration was performed to correct for multiple scattering effects, which can cause underestimations of the OD measurements by microplate readers [17], and to compare OD measurements between the BioTek Synergy H1 microplate reader and the ThermoFisher Genesys spectrophotometer (used as reference).

An overnight culture of *E. coli* K-12 BW25113 grown in M9 medium with 0.9 g/L D-glucose was centrifuged and resuspended in phosphate-buffered saline (PBS) to achieve an initial OD of approximately 4.3. This solution was then serially diluted in PBS to create nine additional solutions with varying cell concentrations. Each solution (1 mL) was added to three wells of a 48-well microplate, and OD (600 nm) measurements were taken using the BioTek Synergy H1 microplate reader. For spectrophotometer measurements, each solution was further diluted in PBS to bring the final OD within the instrument’s linear range (OD 0.01-0.5).

Three cuvettes per solution were measured at 600 nm using the ThermoFisher Genesys spectrophotometer. Calibration was performed by comparing the microplate reader measurements to the spectrophotometer measurements, adjusting for the dilution factor.

### Parameter inference on the auxotrophs growth curves and symbolic regression

The data were baseline-subtracted and corrected for multiple scattering using an interpolation algorithm. The resulting data were then smoothed using a rolling average of 10 time points. We applied a linear change point detection algorithm to the specific growth rate (evaluated with a sliding window of 10 points) for each curve, searching for a single change point within a 16-point window. The curves were subsequently fitted using the logistic model (Eq. 2) and exponential HPM (Eq. 4), with relative error as the loss function (Eq 9). Model selection for each segment was conducted using the corrected AIC score (Eq. 11), with a penalty parameter *β* = 2.

Symbolic regression was performed with Kinbiont’s function `downstream_symbolic_regression`, adapted from the implementation of symbolic regression for Julia in Ref. [11]. We used the binary operations +, *−, ×*, and */*, no unary operations, and all other parameters set to default.

### Decision tree on the ecotoxicological dataset

The dataset of Ref. [16] comprises 15120 growth curves profiling the effects of every possible combination of eight commonly used pollutants across 12 bacterial isolates and a mixture of some of them, as described in the main text. Each chemical was either present at a concentration of 0.1 mg l^*−*1^ or absent [16].

After background subtraction, we fitted the entire dataset using the Richards model (Eq. 6), minimizing the relative error in the loss function. A total of 1367 curves with an average relative error greater than 5% were excluded from further analysis (see Fig. S9). Subsequent analyses were performed on the remaining 13753 growth curves, focusing on the duration of the lag phase, the exponential growth rate, and the total growth inferred from the data.

For each of these observables, we performed decision tree analysis using Kinbiont’s `downstream_decision_ tree_regression`, adapted from the implementation in Ref. [39], with default parameters. The decision tree algorithm employed splitting criteria based on the Gini impurity for the mean of the distributions.

To evaluate the model’s performance, we tested different tree depths and conducted a 10-fold cross-validation for each case (Figs. S12 - S14), randomly splitting the dataset into training and test sets. We did not exclude entire conditions from sampling, as the replicated data were outcomes from biological replicates and separate experiments.

### Synthetic data generation and simulations of deterministic and stochastic microbial dynamics

Kinbiont generates synthetic data by solving selected ODE models using the SCiML solver [40]. As a microbial dynamics simulator, Kinbiont supports both deterministic and stochastic cases, enabling hypothesis testing and precise synthetic benchmarking for each model.

For deterministic simulations, users can choose any ODE model implemented in Kinbiont and specify the preferred numerical integrator. Kinbiont is compatible with all numerical integrators available in SCiML, with the Tsitouras 5*/*4 Runge-Kutta method from `DifferentialEquation.jl` [40] set as the default.

For stochastic simulations, Kinbiont models cell division as a discrete-time Poisson process. The simulation time span is divided into regular intervals of Δ*t*, and at each time step, the population size is updated using the following discrete model:

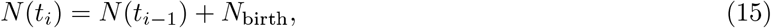

where *N* (*t*_*i*_) represents the number of cells at time *t*_*i*_, and *N*_birth_ denotes the number of birth events in Δ*t*, sampled from a Poisson distribution with rate *N* (*t*_*i−*1_)*µ*Δ*t*:

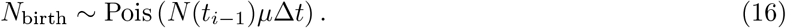

In this context, the growth rate of each individual in the population (*µ*) is calculated based on the concentration of the limiting nutrient, with the user specifying the initial nutrient amount and the culture volume. Various kinetic growth models can be used for this evaluation (see [41] for a review):

1.Monod:

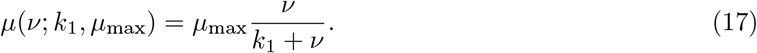

2.Haldane:

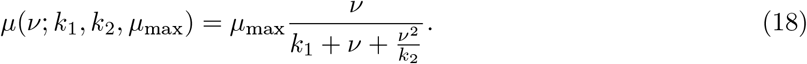

3.Blackman:

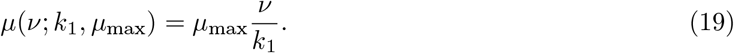

4.Tesseir:

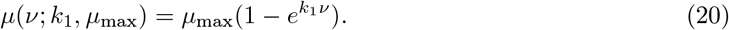

5.Moser:

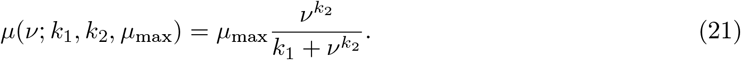

6.Aiba-Edwards:

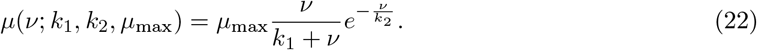

7.Verhulst:

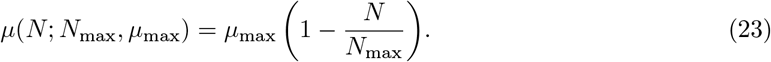

We use *ν* to represent the limiting nutrient concentration throughout, *µ*_max_ denotes the maximum possible growth rate, *k*_1_ (for *i* = 1, 2) is a numerical constant whose specific meaning depends on the model, *N* indicates the number of present cells, and *N*_max_ is the carrying capacity in the Verhulst model.

## Data availability

Source data and Julia scripts to reproduce all the figures are provided in the GitHub repository https://github.com/pinheiroGroup/Kinbiont_utilities.

## Code availability

The entire Kinbiont package is open source and freely available at https://github.com/pinheiroGroup/Kinbiont.jl or via the Julia package manager.

## Acknowledgements

We thank José Davila Velderrain for comments on the manuscript, Claudio Del Fatti for input on early versions of the pipeline, and Martin Ackermann and Gabriele Micali for providing the auxotroph strains. Elements of Figure 1 were produced with BioRender™. This work was funded by Human Technopole.

## Author contributions statement

F.A. and F.P. conceived the method. F.A. implemented the method with input by E.Z.A and F.P.. F.A. generated numerical results. A.P. and F.P. designed experiments. A.P. generated all the experimental data. F.A. and F.P. interpreted the results, generated the figures, and wrote the paper. All authors tested the pipeline. All authors reviewed the manuscript.

## Extended Data Figures

**Extended Data Figure 1:**
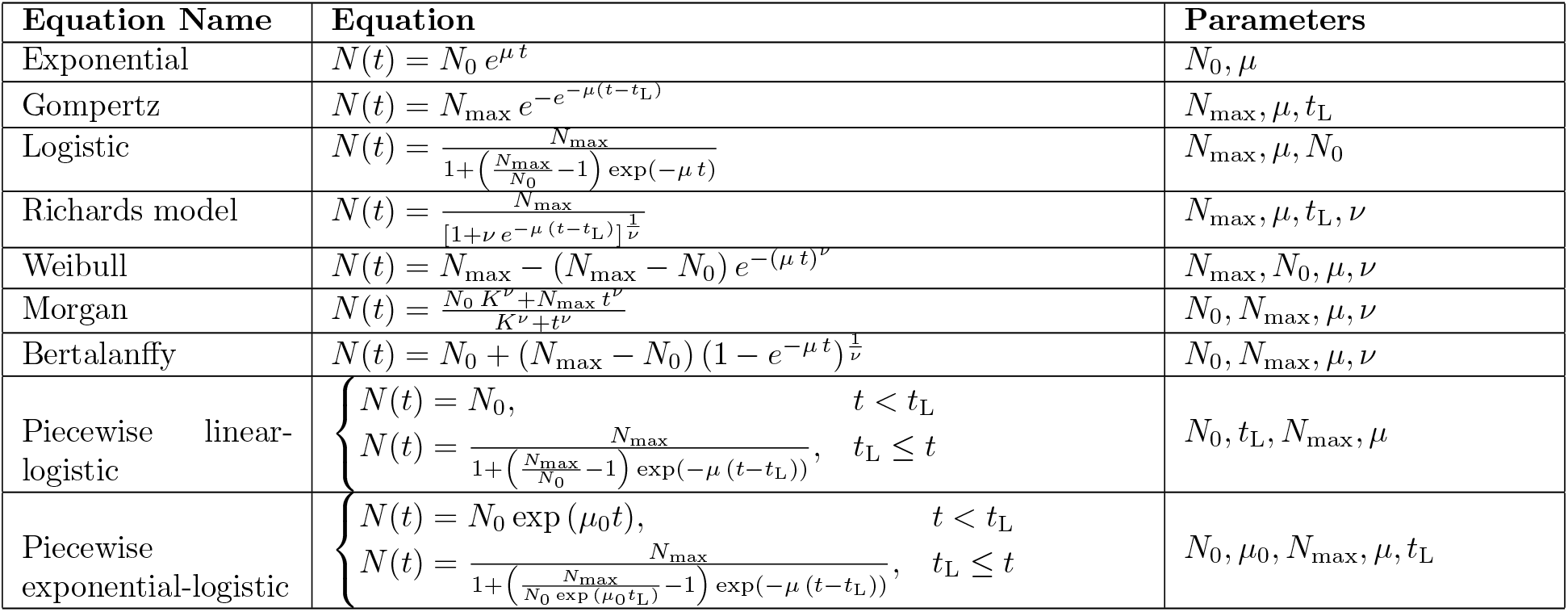
Microbial dynamics models in closed-form solution. List of NL models hard-coded in Kinbiont. While the interpretation of model parameters may differ across models, *N*_0_ refers to the initial population size, *µ* to the exponential growth rate, *N*_max_ to the saturation level or total population growth, *ν* is typically an adjustable shape parameter, and *t*_L_ refers to the lag time [33]. A detailed description of all model-specific parameters is provided in the Supplementary Information.

**Extended Data Figure 2:**
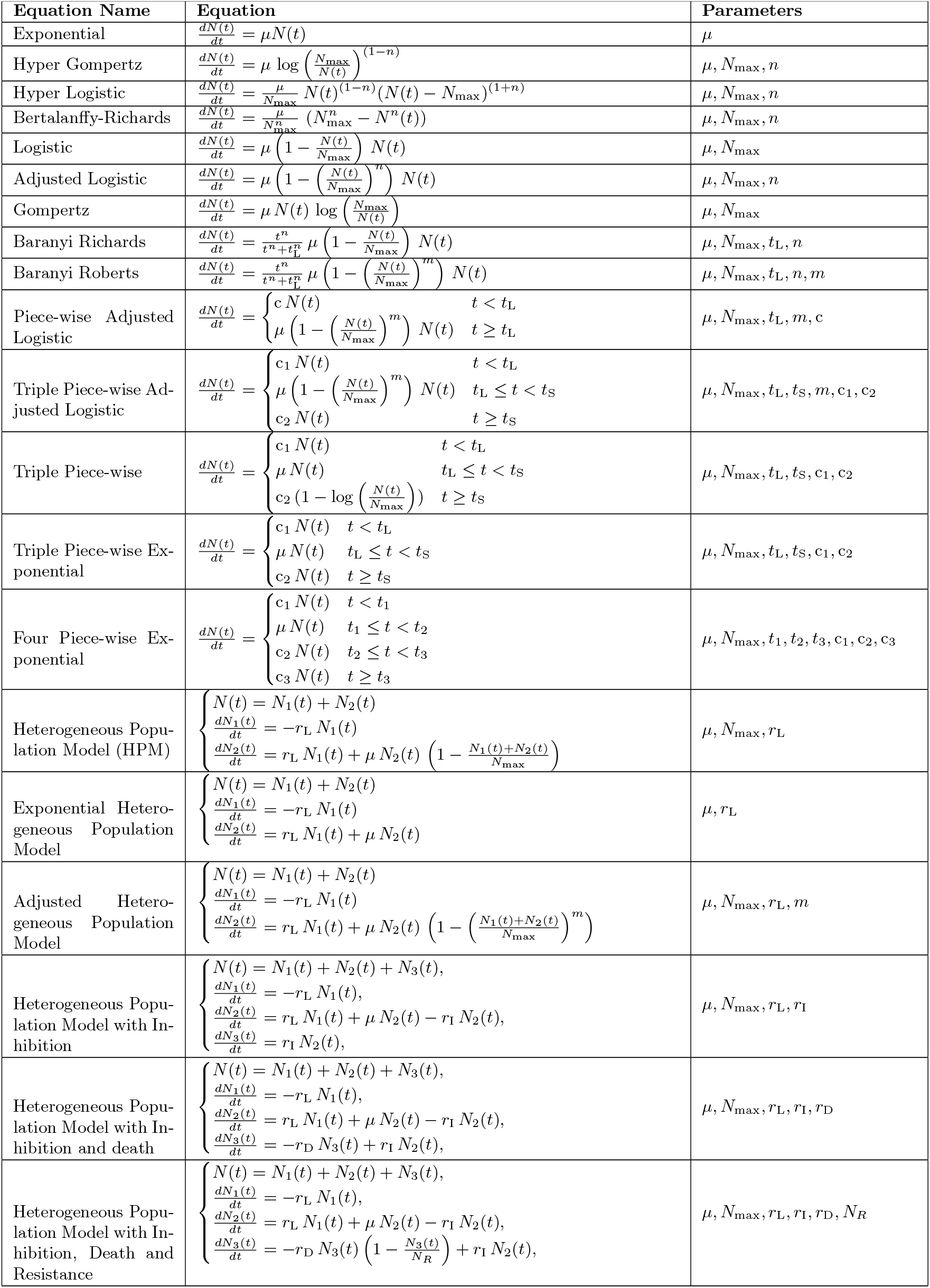
ODE models for microbial dynamics. List of ODE models hard-coded in Kinbiont. While the interpretation of model parameters may vary across models, *N*_0_ refers to the initial population size, *µ* to the exponential growth rate, *N*_max_ to the saturation level or total population growth, *n* and *m* are typically adjustment parameters, and *t*_L_ refers to the lag time. In piece-wise models, *c* and *c*_*i*_ (*i* = 1, …, 3) denote different constants with varying interpretations (e.g., a growth rate in the lag phase), and *t*_*i*_ (*i* = 1, …, 3) denote different time intervals. Additionally, in HPM models, *r*_L_ is the lag rate, *r*_I_ is the inhibition rate, *r*_D_ is the death rate, *N*_*i*_ (*i* = 1, …, 3) refer to the numbers in different sub-populations, and *N*_R_ is the number of resistant cells. A detailed description of all model-specific parameters is provided in the Supplementary Information. 2

**Extended Data Figure 3:**
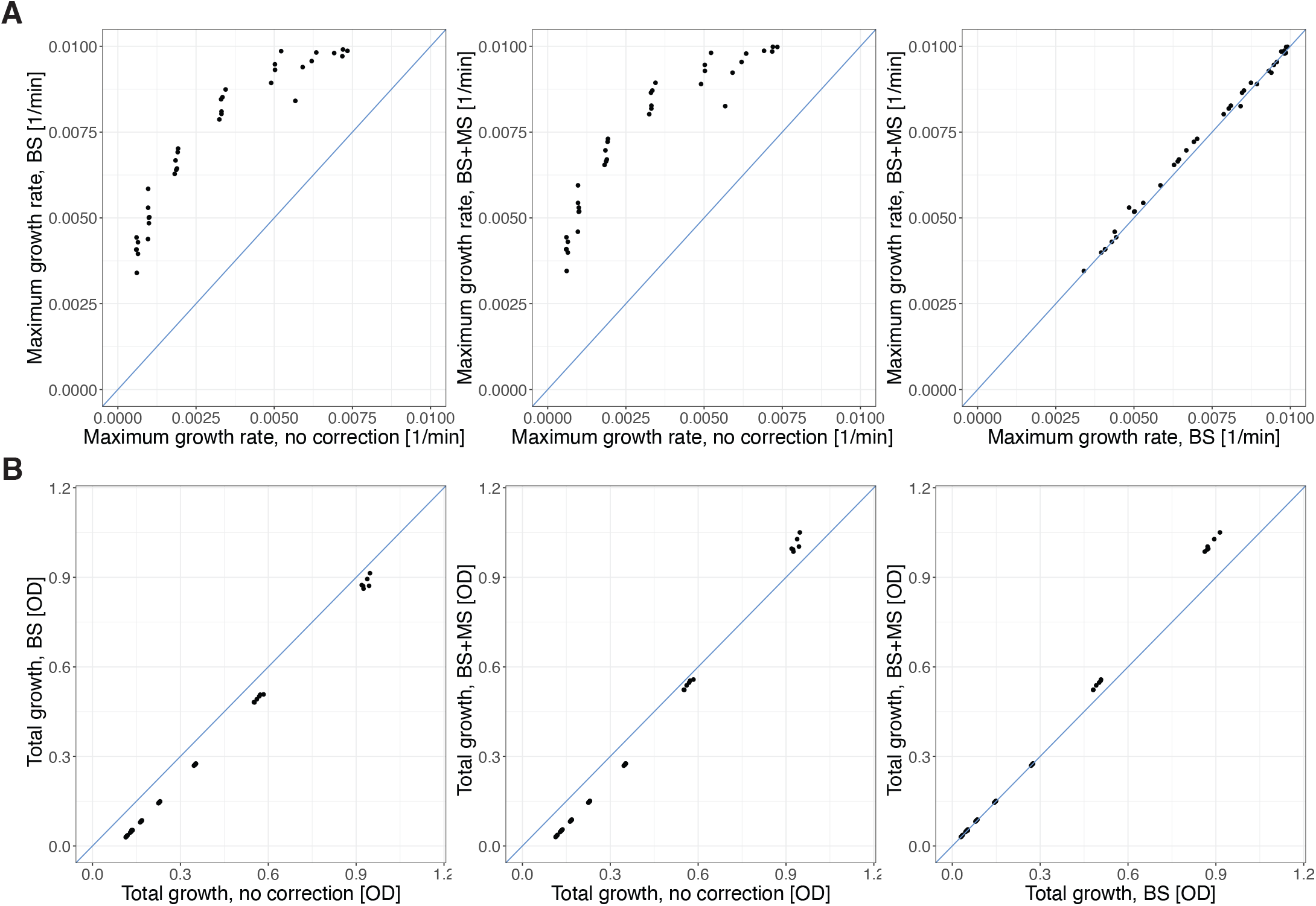
Effects of multiple scattering and background subtraction corrections on inferred parameters: Scatter plots comparing parameters obtained by fitting the data of methionine auxotrophs following pre-processing with different methods: *i)* no pre-processing, *ii)* blank subtraction (BS) using the average value of the blank wells, and *iii)* blank subtraction followed by correction for multiple scattering effects (BS+MS). The solid blue line represents the identity line. **A**, The maximum growth rate estimates are significantly influenced by the background value. In particular, failure to account for background effects leads to an underestimation of growth rates. In contrast, neglecting multiple scattering correction has minimal impact, as the exponential phase typically occurs at OD values within the linear detection range [17]. **B**, The total growth estimate suffers from a small but systematic overestimation due to the absence of blank subtraction. In addition, it is affected by multiple scattering correction, particularly at OD values beyond the linear detection range, where values are underestimated.

**Extended Data Figure 4:**
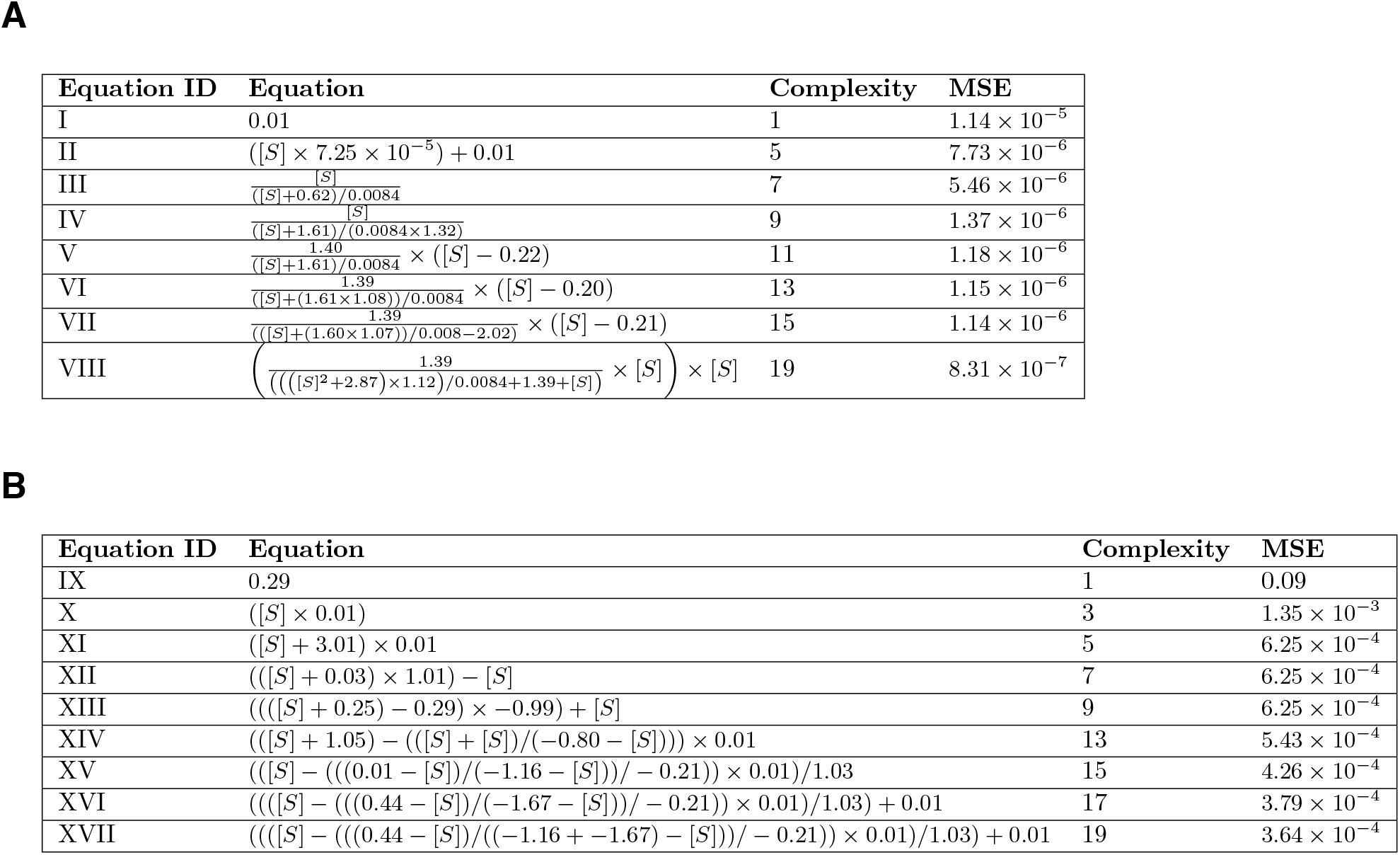
Automatic detection of empirical laws. Hall of Fame of symbolic regression for the growth rate (**A**) and total growth (**B**) for an *E. coli* Δ*metA* knockout expressing a green fluorescent marker (GFP). The model candidates representing the response of these observables as a function of substrate concentration [*S*] are shown together with the mean squared error and the complexity of each proposed law. Equations III and X recapitulate Monod’s empirical laws and are depicted in Fig. 4 along with the measured responses of the different observables.

**Extended Data Figure 5:**
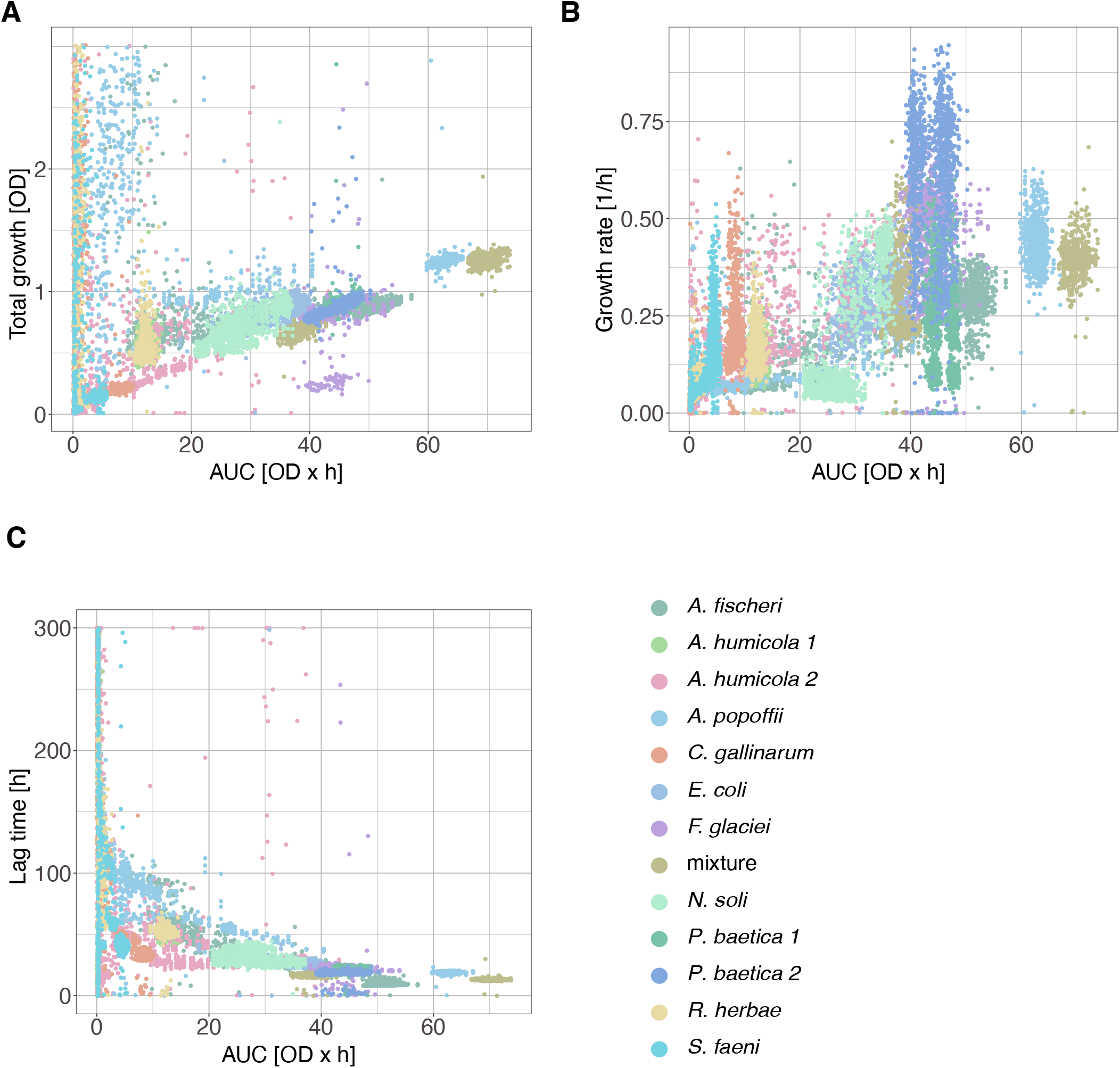
AUC measure vs. model-based fit parameters. AUC vs. the total growth (**A**), growth rate (**B**), and lag time (**C**) inferred with the Richards model (Eq. 6) for all growth curves of Ref. [16]. These analyses suggest that while AUC may serve as a proxy for total growth, it should not be considered a reliable summary variable for growth, as the same AUC value can correspond to significantly different growth rates and lag times. AUC values used in these analyses were taken from Ref. [16].

