## Supplementary Information for "Kinbiont: From time series to ecological and evolutionary responses in microbial systems"

#### List of models implemented in Kinbiont

A comprehensive list of models hard-coded in Kinbiont is given below, together with a description of all model parameters (see also Extended data Figs. 1 and 2).

##### Non-linear function models

- Exponential:

$$N(t) = N_0\, e^{\mu\, t}, \tag{1}$$

where $\mu$ is the growth rate, and $N_0$ it is the initial population size.

- Gompertz:

$$N(t) = N_{\max}\, e^{-e^{-\mu\,(t-t_{\mathrm{L}})}}, \tag{2}$$

where $\mu$ is the growth rate, $N_{\max}$ is the total growth, and $t_{\mathrm{L}}$ the lag time.

- Logistic:

$$N(t) = \frac{N_{\max}}{1+\left(\frac{N_{\max}}{N_0}-1\right)\exp\left(-\mu\, t\right)}, \tag{3}$$

where $\mu$ is the growth rate, $N_0$ it is the initial population size, and $N_{\max}$ the total growth.

*Correspondence to

- Richards model:
$$N(t) = \frac{N_{\max}}{\left[1 + \nu\, e^{-\mu\,(t - t_{\mathrm{L}})}\right]^{\frac{1}{\nu}}}, \tag{4}$$
where $\mu$ is the growth rate, $N_{\max}$ is the total growth, $t_{\mathrm{L}}$ the lag time, and $\nu$ a shape constant.

- Weibull:
$$N(t) = N_{\max} - (N_{\max} - N_0)\, e^{-(\mu\, t)^{\nu}}, \tag{5}$$
where $\mu$ is the growth rate, $N_0$ it is the initial population size, $N_{\max}$ the total growth and $\nu$ a shape constant.

- Morgan:
$$N(t) = \frac{N_0\, K^{\nu} + N_{\max}\, t^{\nu}}{K^{\nu} + t^{\nu}}, \tag{6}$$
$N_0$ it is the initial population size, $N_{\max}$ the total growth, $K$ is the time where half-maximal growth is achieved, and $\nu$ a shape constant.

- Bertalanffy:
$$N(t) = N_0 + (N_{\max} - N_0)\,(1 - e^{-\mu\, t})^{\frac{1}{\nu}}, \tag{7}$$
$N_0$ is the starting condition, $N_{\max}$ the total growth, $\mu$ the growth rate, and $\nu$ a shape constant.

- Piece-wise Linear-Logistic
$$\begin{cases} N(t) = N_0, & t < t_{\mathrm{L}} \\ N(t) = \dfrac{N_{\max}}{1 + \left(\frac{N_{\max}}{N_0} - 1\right) \exp\left(-\mu\,(t - t_{\mathrm{L}})\right)} & t_{\mathrm{L}} \leq t, \end{cases} \tag{8}$$
$N_0$ is the starting condition, $N_{\max}$ the total growth, $\mu$ the growth rate, and $t_{\mathrm{L}}$ the lag time.

- Piece-wise Exponential-Logistic:
$$\begin{cases} N(t) = N_0 \exp\left(\mu_0 t\right), & t < t_{\mathrm{L}} \\ N(t) = \dfrac{N_{\max}}{1 + \left(\frac{N_{\max}}{N_0 \exp\left(\mu_0 t_{\mathrm{L}}\right)} - 1\right) \exp\left(-\mu\,(t - t_{\mathrm{L}})\right)} & t_{\mathrm{L}} \leq t, \end{cases} \tag{9}$$
$N_0$ is the starting condition, $N_{\max}$ the total growth, $\mu$ the growth rate, $t_{\mathrm{L}}$ the lag time, and $\mu_0$ the growth during the lag phase.

#### Ordinary differential equation models

- Exponential:

$$\frac{dN(t)}{dt} = \mu N(t), \tag{10}$$

where $\mu$ is the growth rate.

- Hyper Gompertz [1]:

$$\frac{dN(t)}{dt} = \mu \log \left( \frac{N_{\text{max}}}{N(t)} \right)^{(1-n)}, \tag{11}$$

where $\mu$ is the growth rate, $N_{\text{max}}$ the total growth and $n$ a shape constant.

- Hyper Logistic [1]:

$$\frac{dN(t)}{dt} = \frac{\mu}{N_{\text{max}}} N(t)^{(1-n)} \left( N(t) - N_{\text{max}} \right)^{(1+n)}, \tag{12}$$

where $\mu$ is the growth rate, $N_{\text{max}}$ the total growth and $n$ a shape constant.

- Bertalanffy-Richards:

$$\frac{dN(t)}{dt} = \frac{\mu}{N_{\text{max}}^n} \left( N_{\text{max}}^n - N^n(t) \right), \tag{13}$$

where $\mu$ is the growth rate, $N_{\text{max}}$ the total growth, and $n$ a shape constant.

- Logistic [1]:

$$\frac{dN(t)}{dt} = \mu \left( 1 - \frac{N(t)}{N_{\text{max}}} \right) N(t), \tag{14}$$

where $\mu$ is the growth rate, and $N_{\text{max}}$ the total growth.

- Adjusted Logistic:

$$\frac{dN(t)}{dt} = \mu \left( 1 - \left( \frac{N(t)}{N_{\text{max}}} \right)^n \right) N(t), \tag{15}$$

where $\mu$ is the growth rate, $N_{\text{max}}$ the total growth and $n$ a shape constant.

- Gompertz [1]:

$$\frac{dN(t)}{dt} = \mu\, N(t)\, \log\left(\frac{N_{\mathrm{max}}}{N(t)}\right), \tag{16}$$

where $\mu$ is the growth rate, and $N_{\mathrm{max}}$ the total growth.

- Baranyi-Richards [2]:

$$\frac{dN(t)}{dt} = \frac{t^n}{t^n + t_{\mathrm{L}}^n}\, \mu \left(1 - \frac{N(t)}{N_{\mathrm{max}}}\right) N(t), \tag{17}$$

where $\mu$ is the growth rate, $N_{\mathrm{max}}$ the total growth, $t_{\mathrm{L}}$ is the lag time and $n$ a shape constant.

- Baranyi-Roberts [3]:

$$\frac{dN(t)}{dt} = \frac{t^n}{t^n + t_{\mathrm{L}}^n}\, \mu \left(1 - \left(\frac{N(t)}{N_{\mathrm{max}}}\right)^m\right) N(t), \tag{18}$$

where $\mu$ is the growth rate, $N_{\mathrm{max}}$ the total growth, $t_{\mathrm{L}}$ is the lag time, $n$ and $m$ are shape constants.

- Piece-wise Adjusted Logistic:

$$\frac{dN(t)}{dt} = \begin{cases} c\, N(t) & \text{for } t < t_{\mathrm{L}} \\ \mu \left(1 - \left(\frac{N(t)}{N_{\mathrm{max}}}\right)^m\right) N(t) & \text{for } t \geq t_{\mathrm{L}}, \end{cases} \tag{19}$$

where $\mu$ is the growth rate, $N_{\mathrm{max}}$ the total growth, $t_{\mathrm{L}}$ is the lag time, $m$ is shape constant, and $c$ the growth rate during the lag phase (it can be 0).

- Triple Piece-wise Adjusted Logistic:

$$\frac{dN(t)}{dt} = \begin{cases} c_1\, N(t) & \text{for } t < t_{\mathrm{L}} \\ \mu \left(1 - \left(\frac{N(t)}{N_{\mathrm{max}}}\right)^m\right) N(t) & \text{for } t_{\mathrm{L}} \leq t < t_{\mathrm{S}} \\ c_2\, N(t) & \text{for } t \geq t_{\mathrm{S}}, \end{cases} \tag{20}$$

where $\mu$ is the growth rate, $N_{\mathrm{max}}$ the total growth, $t_{\mathrm{L}}$ is the lag time, $m$ is a shape constant, $c_1$ the growth rate during the lag phase (it can be 0), $t_{\mathrm{S}}$ the time when the saturation phase starts, and $c_2$

the growth rate after the theoretical saturation point (e.g., to account for a linear increase in the time series when the growth curve should in principle be in the stationary phase).

- Triple Piece-wise:

$$\frac{dN(t)}{dt} = \begin{cases} c_1\, N(t) & \text{for } t < t_{\mathrm{L}} \\ \mu\, N(t) & \text{for } t_{\mathrm{L}} \le t < t_{\mathrm{S}} \\ c_2 \left(1 - \log\left(\frac{N(t)}{N_{\max}}\right)\right) & \text{for } t \ge t_{\mathrm{S}}, \end{cases} \tag{21}$$

where $\mu$ is the growth rate, $N_{\max}$ the total growth, $t_{\mathrm{L}}$ is the lag time, $c_1$ is the growth rate during the lag phase (it can be 0), $t_{\mathrm{S}}$ is the time when stationary phase starts, and $c_2$ is the growth rate during after the theoretical saturation point (e.g., to account for a linear increase in the time series when the growth curve should in principle be in the stationary phase).

- Triple Piece-wise Exponential:

$$\frac{dN(t)}{dt} = \begin{cases} c_1\, N(t) & \text{for } t < t_{\mathrm{L}} \\ \mu\, N(t) & \text{for } t_{\mathrm{L}} \le t < t_{\mathrm{S}} \\ c_2\, N(t) & \text{for } t \ge t_{\mathrm{S}}, \end{cases} \tag{22}$$

where $\mu$ is the growth rate, $N_{\max}$ the total growth, $t_{\mathrm{L}}$ is the lag time, $c_1$ is the growth rate during the lag phase (it can be 0), $t_{\mathrm{S}}$ is the time when stationary phase starts, and $c_2$ is the growth rate during after the theoretical saturation point (e.g., to account for a linear increase in the time series when the growth curve should in principle be in the stationary phase).

- Four Piece-wise Exponential:

$$\frac{dN(t)}{dt} = \begin{cases} c_1\, N(t) & \text{for } t < t_1 \\ \mu\, N(t) & \text{for } t_1 \le t < t_2 \\ c_2\, N(t) & \text{for } t_2 \le t < t_3 \\ c_3\, N(t) & \text{for } t \ge t_3, \end{cases} \tag{23}$$

where $\mu$ is the growth rate, $N_{\max}$ is the maximum population size, $t_1$ is the lag time, $c_1$ is the growth rate during the lag phase (which can be 0), $t_2$ is the time when growth occurs after the exponential phase, $c_2$ is the growth rate during this phase, $t_3$ marks the start of the stationary phase, and $c_3$ is the growth rate during the stationary phase.

- Heterogeneous Population Model (HPM) [4]:

$$\begin{cases} N(t) = N_1(t) + N_2(t) \\ \frac{dN_1(t)}{dt} = -r_{\mathrm{L}}\, N_1(t) \\ \frac{dN_2(t)}{dt} = r_{\mathrm{L}}\, N_1(t) + \mu\, N_2(t)\left(1 - \dfrac{N_1(t) + N_2(t)}{N_{\max}}\right), \end{cases} \tag{24}$$

where $N_1$ is the population of dormant cells, $N_2$ is the population of active cells capable of duplicating, $\mu$ is the growth rate, $N_{\max}$ is the total growth, and $r_{\mathrm{L}}$ is the lag rate, defined as the rate of transition between the $N_1$ and $N_2$ populations. Here, we assume that all cells are in the dormant state at the start (i.e., $N_1(t = 0) = \mathrm{OD}(t = 0)$, and $N_2(t = 0) = 0$).

- Exponential Heterogeneous Population Model:

$$\begin{cases} N(t) = N_1(t) + N_2(t) \\ \frac{dN_1(t)}{dt} = -r_{\mathrm{L}}\, N_1(t) \\ \frac{dN_2(t)}{dt} = r_{\mathrm{L}}\, N_1(t) + \mu\, N_2(t), \end{cases} \tag{25}$$

where similarly to the HPM model, $N_1$ and $N_2$ refer to the populations of dormant and active cells, respectively. $\mu$ is the growth rate, and the lag rate $r_{\mathrm{L}}$ denotes the transition between the $N_1$ and $N_2$ populations. Here, we also assume that all cells are in the dormant state at the start (i.e., $N_1(t = 0) = \mathrm{OD}(t = 0)$, and $N_2(t = 0) = 0$).

- Adjusted Heterogeneous Population Model:

$$\begin{cases} N(t) = N_1(t) + N_2(t) \\ \frac{dN_1(t)}{dt} = -r_{\mathrm{L}}\, N_1(t) \\ \frac{dN_2(t)}{dt} = r_{\mathrm{L}}\, N_1(t) + \mu\, N_2(t)\left(1 - \left(\dfrac{N_1(t) + N_2(t)}{N_{\max}}\right)^{m}\right), \end{cases} \tag{26}$$

where similarly to all variants of the HPM model, $N_1$ and $N_2$ refer to the populations of dormant and active cells, respectively. $\mu$ is the growth rate, $N_{\max}$ the total growth, $t_{\mathrm{L}}$ is the lag rate (i.e. the rate of transition between $N_1(t)$ and $N_2(t)$) and $m$ a shape constant. Here, we also assume that all cells are in the dormant state at the start (i.e., $N_1(t=0) = \mathrm{OD}(t=0)$, and $N_2(t=0) = 0$).

- Heterogeneous Population Model with Inhibition:

$$\begin{cases} N(t) = N_1(t) + N_2(t) + N_3(t) \\ \frac{dN_1(t)}{dt} = -r_{\mathrm{L}}\, N_1(t) \\ \frac{dN_2(t)}{dt} = r_{\mathrm{L}}\, N_1(t) + \mu\, N_2(t) - r_{\mathrm{I}}\, N_2(t) \\ \frac{dN_3(t)}{dt} = r_{\mathrm{I}}\, N_2(t), \end{cases} \tag{27}$$

where $N_1$, $N_2$, and $N_3$ refer to the populations of dormant, active, and inhibited cells. In contrast to the population of actively duplicating cells in $N_2$, cells in the $N_3$ inhibited state are active but do not duplicate. The other parameters are the growth rate $\mu$, the lag rate for the transition between $N_1$ and $N_2$, $r_{\mathrm{L}}$, and the lag rate for the transition between $N_2$ and $N_3$, $r_{\mathrm{I}}$. As in the cases above, we assume that all cells start in the dormant state (i.e., $N_1(t=0) = \mathrm{OD}(t=0)$, $N_2(t=0) = N_3(t=0) = 0$).

- Heterogeneous Population Model with Inhibition and Death:

$$\begin{cases} N(t) = N_1(t) + N_2(t) + N_3(t) \\ \frac{dN_1(t)}{dt} = -r_{\mathrm{L}}\, N_1(t) \\ \frac{dN_2(t)}{dt} = r_{\mathrm{L}}\, N_1(t) + \mu\, N_2(t) - r_{\mathrm{I}}\, N_2(t) \\ \frac{dN_3(t)}{dt} = -r_{\mathrm{D}}\, N_3(t) + r_{\mathrm{I}}\, N_2(t), \end{cases} \tag{28}$$

where $N_1$, $N_2$, and $N_3$ refer to the populations of dormant, active, and inhibited cells. In contrast to the population of actively duplicating cells in $N_2$, cells in the $N_3$ inhibited state are active but do not duplicate. The other parameters are the growth rate $\mu$, the lag rate for the transition between $N_1$ and $N_2$, $r_{\mathrm{L}}$, the lag rate for the transition between $N_2$ and $N_3$, $r_{\mathrm{I}}$, and the rate at which cells die, $r_{\mathrm{D}}$. As in the cases above, we assume that all cells start in the dormant state (i.e., $N_1(t=0) = \mathrm{OD}(t=0)$, $N_2(t=0) = N_3(t=0) = 0$).

- Heterogeneous Population Model with Inhibition, Death and Resistance:

$$
\begin{cases}
N(t) = N_1(t) + N_2(t) + N_3(t) \\
\frac{dN_1(t)}{dt} = -r_{\mathrm{L}}\, N_1(t) \\
\frac{dN_2(t)}{dt} = r_{\mathrm{L}}\, N_1(t) + \mu\, N_2(t) - r_{\mathrm{I}}\, N_2(t) \\
\frac{dN_3(t)}{dt} = -r_{\mathrm{D}}\, N_3(t) \left(1 - \dfrac{N_3(t)}{N_{\mathrm{R}}}\right) + r_{\mathrm{I}}\, N_2(t),
\end{cases}
\tag{29}
$$

where $\mu$ is the growth rate, $t_{\mathrm{L}}$ is the lag rate (i.e. the rate of transition between $N_1(t)$ and $N_2(t)$), $r_{\mathrm{I}}$ is the rate of which cell are inhibited (i.e. the rate of transition between $N_2(t)$ and $N_3(t)$), $r_{\mathrm{D}}$ is the rate of which cell are die, and $N_{\mathrm{R}}$ it the number of cell that will become inactive but do not die.

### List of error functions

Kinbiont includes the following expressions for the loss function ($\mathcal{L}(P)$ where $P$ is a set of model specific parameters):

1. $L2$ norm of the difference between the numerical solution (i.e., $\hat{N}(t_i, P)$ for $i = 1, \dots, n$, where $n$ is the number of data points) of the desired model and the given data (i.e., $N(t_i)$ for $i = 1, \dots, n$):

$$
\mathcal{L}(P) = \frac{1}{n} \sum_{i=1}^{n} \left(N(t_i) - \hat{N}(t_i, P)\right)^2, \tag{30}
$$

where $n$ is the number of data points.

2. Relative error between the solution and data:

$$
\mathcal{L}(P) = \frac{1}{n} \sum_{i=1}^{n} \frac{1}{2} \left(1 - \frac{N(t_i)}{\hat{N}(t_i, P)}\right)^2, \tag{31}
$$

where $n$ is the number of data points.

3. $L2$ norm of the difference between the specific growth rate of the numerical solution of the desired model and the corresponding derivatives of the data:

$$
\mathcal{L}(P) = \frac{1}{n} \sum_{i=1}^{n} \left(\frac{dN(t_i)}{dt} - \frac{d\hat{N}(t_i, P)}{dt}\right)^2, \tag{32}
$$

where $n$ is the number of data points.

4. Logarithm of the $L2$ norm of the difference between the numerical solution of the desired model and the given data:

$$\mathcal{L}(P) = \log\left(\frac{1}{n}\sum_{i=1}^{n}\left(N(t_i) - \hat{N}(t_i, P)\right)^2\right), \tag{33}$$

   where $n$ is the number of data points.

5. Logarithm of the relative error between the solution and data:

$$\mathcal{L}(P) = \log\left(\frac{1}{n}\sum_{i=1}^{n}\frac{1}{2}\left(1 - \frac{N(t_i)}{\hat{N}(t_i, P)}\right)^2\right), \tag{34}$$

   where $n$ is the number of data points.

6. $L2$ norm of the difference between the numerical solution of the desired model and the data, weighted by the standard deviation of empirical blank data:

$$\mathcal{L}(P) = \frac{1}{n}\sum_{i=1}^{n}\left(\frac{N(t_i) - \hat{N}(t_i, P)}{\text{std blank}}\right)^2, \tag{35}$$

   where std blank is the standard deviation of the empirical blank data, and $n$ is the number of data points.

**Comparison of parameter inference between ODE and NL fit**

To compare the performance of the nonlinear (NL) and ordinary differential equation (ODE) fitting schemes, we carried out an analysis using synthetic data. We used the logistic model, which is a default option in Kinbiont for both fitting methods; the explicit expressions are provided in the row corresponding to the logistic model in Extended Data Figs. 1 and 2 (respectively the third and the fifth row) . Importantly, the NL version of the model includes an additional parameter, as the initial value of the function is not constrained to the observed measurement as it is in the ODE fits.

Using Kinbiont, we generated 100 synthetic logistic curves with random parameters sampled from uniform distributions: $\mu \sim \mathcal{U}(0.01, 0.1)$, $N_{\max} \sim \mathcal{U}(0.3, 2.5)$, and $N_0 \sim \mathcal{U}(0.1, N_{\max}/2)$. Gaussian noise with a mean of 0 and $2\sigma = \{0.01, 0.03, 0.05, 0.07\}$ was added to each point of the curves. The resulting curves were then fit using the corresponding ODE and NL methods. The loss function (average relative error) and inferred model parameters are presented in Figs. S1 and S2.

As shown in Fig. S1, the ODE model consistently yields a lower loss function when estimating parameters. Differences in parameter estimates, likely due to the additional degree of freedom in the NL model, can affect the stability of the results. However, these differences can be mitigated by adjusting the optimization scheme, such as increasing the number of iterations, making more informed initial parameter guesses or using a multistart approach. Fig. S2 further corroborates these findings by comparing the inferred parameters to the ground truth, demonstrating that both methods produce accurate inferences of microbial dynamics parameters. These results also emphasize the reliability of ODE model fits, particularly in scenarios where closed-form solutions for NL models are unavailable.

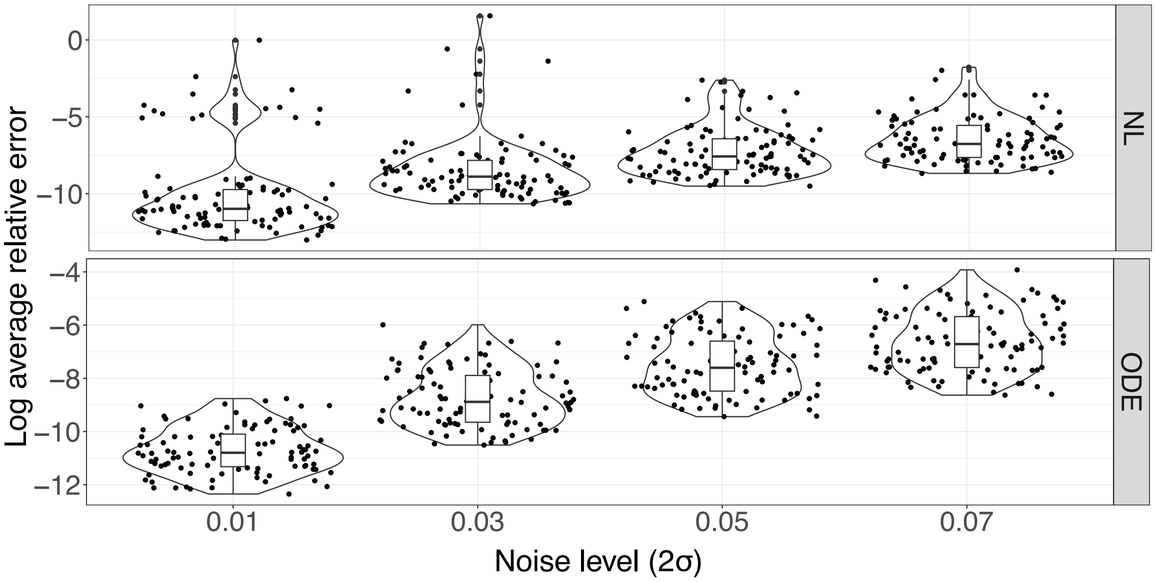


Supplementary Figure 1: **Distributions of average relative error for NL and ODE logistic model fits:** Distribution of the logarithm of the average relative error obtained by fitting 100 synthetic logistic curves with four different levels of Gaussian noise using the NL (upper panel) and ODE (lower panel) fitting schemes. While both methods perform well on synthetic data with noise, fitting the model in ODE form consistently yields lower average loss function values. Differences in parameter estimates provided by these two methods should not affect biological conclusions (see Fig. S2) and, as discussed in the text, can be mitigated by adjusting fit parameters (e.g., the chosen optimizer, number of iterations, tolerance, stopping criteria, or improved initial condition guesses).

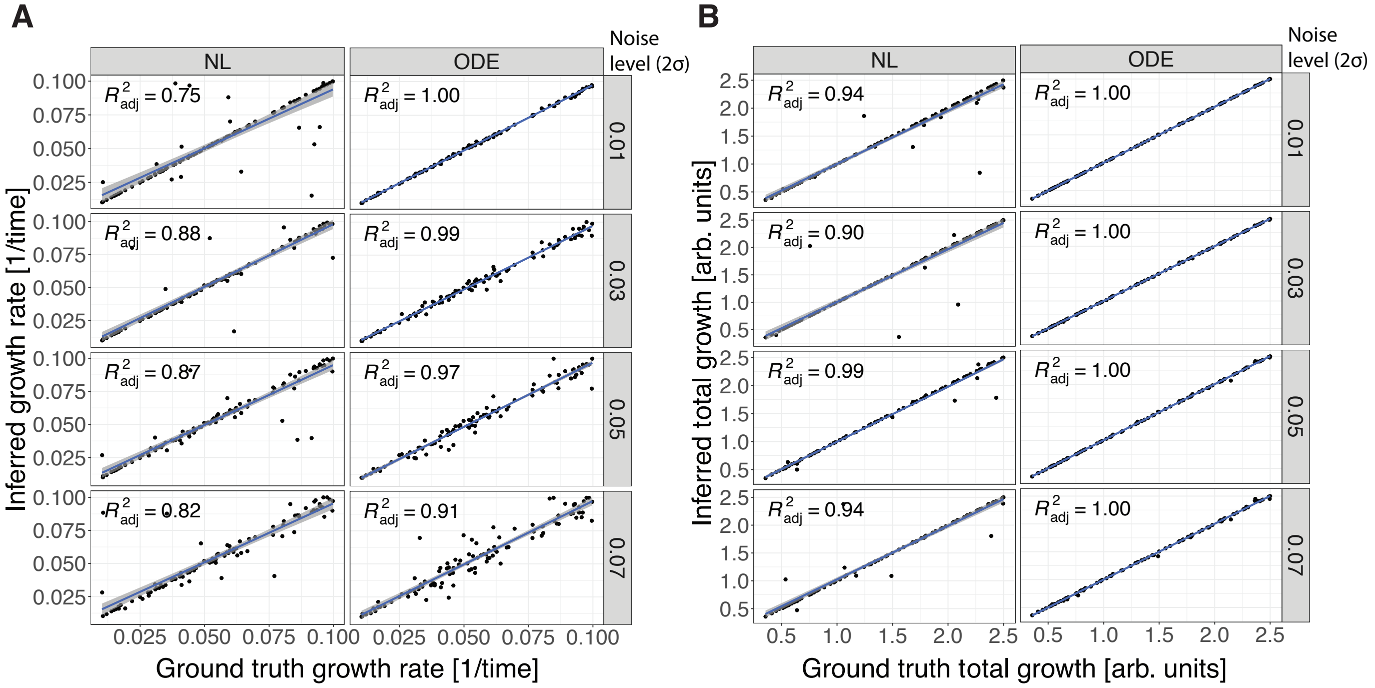


Supplementary Figure 2: **Parameters inference benchmark:** Scatter plots comparing the ground truth parameters with those inferred from fitting 100 synthetic logistic curves using NL and ODE fitting schemes under four different levels of Gaussian noise. **A**, Logistic growth parameter $\mu$. **B**, Total growth $N_{\max}$. Each plot includes a linear fit (blue line) with a shaded 95% confidence interval (gray) and the adjusted $R^2$ correlation coefficient is annotated.

##### Comparison of the detection of the exponential growth rate using a model-based or a non-parametric method

Given that different models interpret the growth rate $\mu$ differently, direct comparisons of $\mu$ values across models are not always straightforward [5]. A common practice in bacterial kinetics analysis is to focus on the maximum exponential growth rate, expressed as $\mu_{\max} = \max\left(\frac{d\log(N(t))/N_0}{dt}\right)$, where $N$ is the measurement and $N_0$ is the initial value. In batch culture, $\mu_{\max}$ is usually obtained from the specific growth rate, which is typically determined through a linear fit to the logarithm of the growth curve. Applying this method within a sliding window allows for tracking the dynamics of the exponential growth rate over time and identifying its maximum value or average.

In Fig. S3, we compare the ground truth maximum exponential/specific growth rate with estimates obtained using both an ODE-based approach and the sliding window method with different window sizes. This comparison involved generating 100 synthetic curves using the adjusted HPM model (cf. Extended Data Fig. 2) with parameters sampled from uniform distributions: $\mu \sim \mathcal{U}(0.01, 0.08)$, $N_{\max} \sim \mathcal{U}(0.4, 1.5)$, $n \sim \mathcal{U}(0.07, 5)$, and $r_L \sim \mathcal{U}(0.02, 0.09)$. Gaussian noise with a mean of 0 and $2\sigma = 0.01, 0.03, 0.05, 0.07$ was added to each data point to evaluate the effect of noise on parameter inference accuracy.

The results indicate that the ODE-based approach consistently provides more accurate maximum growth rate estimates than the sliding window method. Since the fitted model is inherently noise-free, this method eliminates the need for hyperparameters, such as the smoothing window size, making it a robust alternative for obtaining maximum growth rates. This approach also enhances the detection and standardization of growth rate observables. For completeness, Kinbiont reports both the growth rates of the selected model and the maximum exponential growth rate evaluated on the growth curve generated by the inferred model.

#### Change point detection inference

To benchmark the accuracy of the signal processing methods integrated in Kinbiont for detecting changes in microbial growth dynamics, we performed an *in silico* test. We generated 1000 time series, each containing a change point, using the following procedure:

First, we simulated the HPM model (cf. Extended Data Fig. 2) with parameters $\mu_{\max} = 0.06$, $r_L = 0.01$, and $N\max = 1.01$ over the time interval $(0, t_{cp})$, where the change point time $t_{cp}$ was drawn from a uniform distribution $t_{cp} \sim \mathcal{U}(150, 450)$. To ensure continuity, we then simulated a second logistic model (cf. Extended Data Fig. 2) with parameters $\mu_{\max} = 0.08$ and $N_{\max} = 2$ over the time span $(t_{cp}, 600)$. Gaussian noise with a mean of 0 and $2\sigma = 0.01, 0.03, 0.05, 0.07$ was added to each generated curve.

We applied two change point detection methods implemented in Kinbiont: the least square density difference (LSDD) method [6] and the sliding window with linear fitting approach (described in the Methods section). In Fig. S4, we compared the inferred change point times with the ground truth values. Both methods demonstrated strong performance and were effective in detecting changes in microbial growth dynamics. However, a systematic error was observed in the change point estimates using the LSDD method due to the implementation in `ChangePointDetection.jl`, where the dissimilarity curve's value is assigned to the start of the window rather than the middle. This deviation can be easily corrected by shifting the estimate by half the window size, according to the user's needs.

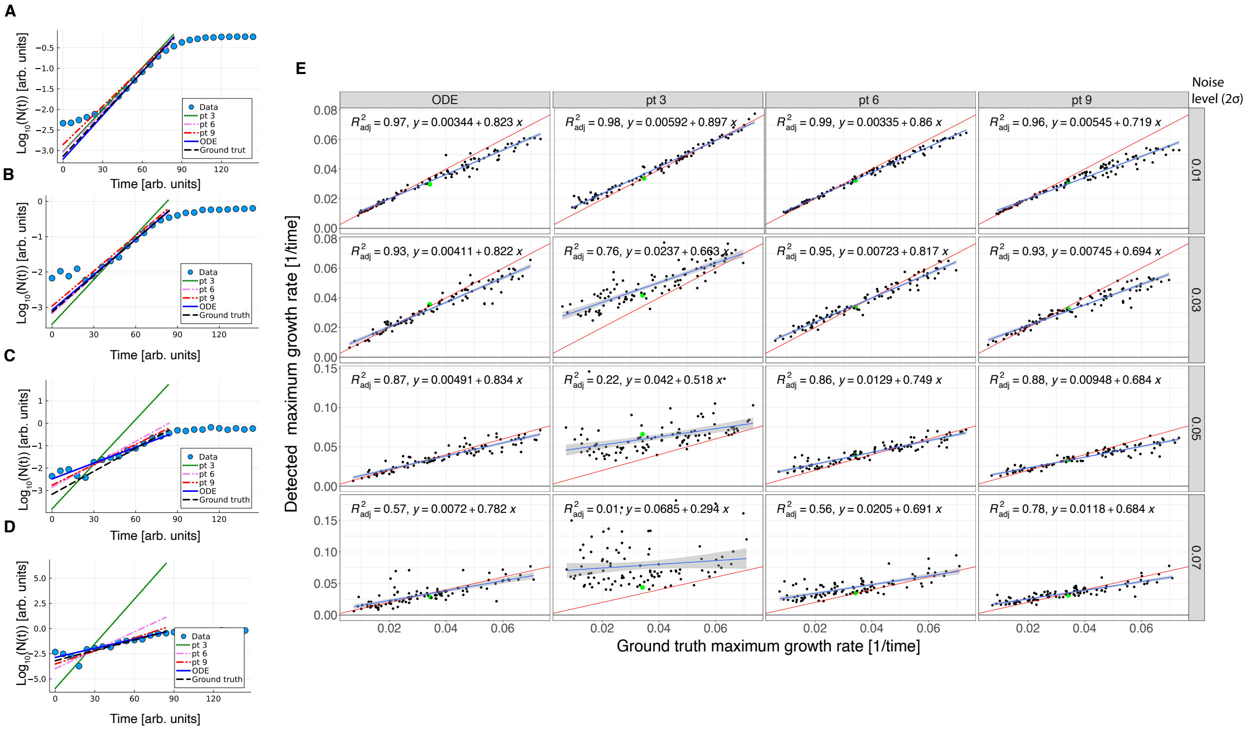

Supplementary Figure 3: **Detection of maximum exponential growth rate:** Comparison of the maximum growth rate estimates obtained using both model-based and non-parametric methods on a synthetic dataset (100 curves) generated with the aHPM model under different levels of Gaussian noise. The synthetic data were used to fit the aHPM model to each curve, and the maximum growth rate was subsequently evaluated using the inferred parameters (ODE) and log-linear fits on a sliding window with varying sizes (3 to 9 points, in steps of 3). **A - D** show examples of the estimated maximum growth rate alongside the data (dots) in a semi-logarithmic scale, illustrating the results from different methods. In **E**, scatter plots compare the inferred values to the ground truth for the maximum exponential growth rate across the 100 synthetic curves. The red line represents the identity line (perfect fit), the blue line shows the linear fit, the gray area indicates the 95% confidence interval, and the green dots correspond to the parameter values for the linear plots in **A - D**. In particular, the linear fits and the correlation coefficient show that ODE fits outperform the sliding window approach in the detection of maximum growth rate at any noise level. While a larger sliding window works better in comparison to a smaller ones at higher noise, it underestimates higher values of the growth rate. When using the latter method, users should set the size of the window according to the noise level and rate of data acquisition.

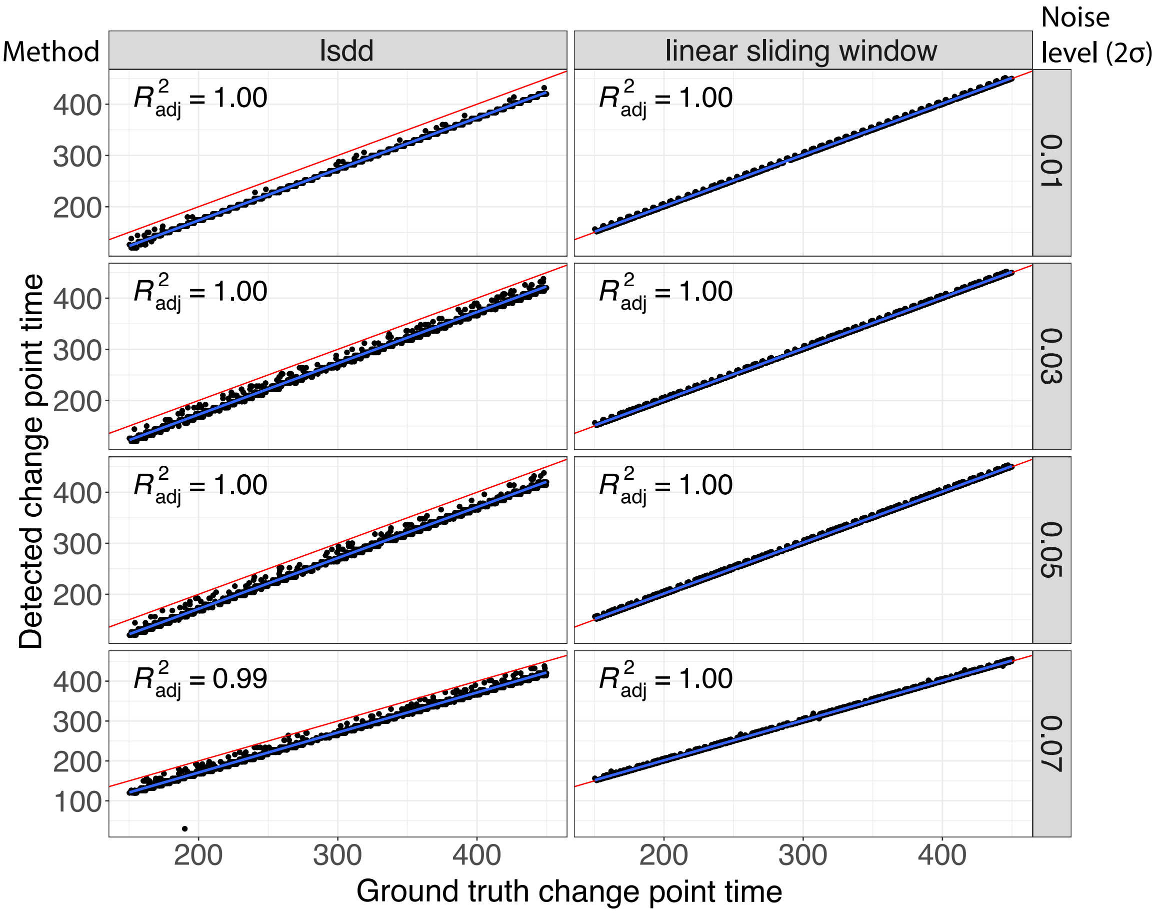


Supplementary Figure 4: **Change point detection benchmark:** Comparison of change point detection algorithms using 1000 synthetic time series. The x-axis represents the ground truth change point, and the y-axis shows the inferred value. Blue lines indicate a linear fit, with gray areas representing the 95% confidence interval, and the adjusted $R^2$ correlation coefficient is noted. Each row corresponds to a different noise level, and the two columns compare the change point detection methods implemented in Kinbiont. As discussed in the Supplementary Information, the systematic difference between the LSDD and linear window methods arises from the choice to assign the value of the dissimilarity curve to the start of the window in the LSDD method, and to the window mid-point when using the linear window method.

#### Supporting figures for the parameter inference of complex microbial kinetics

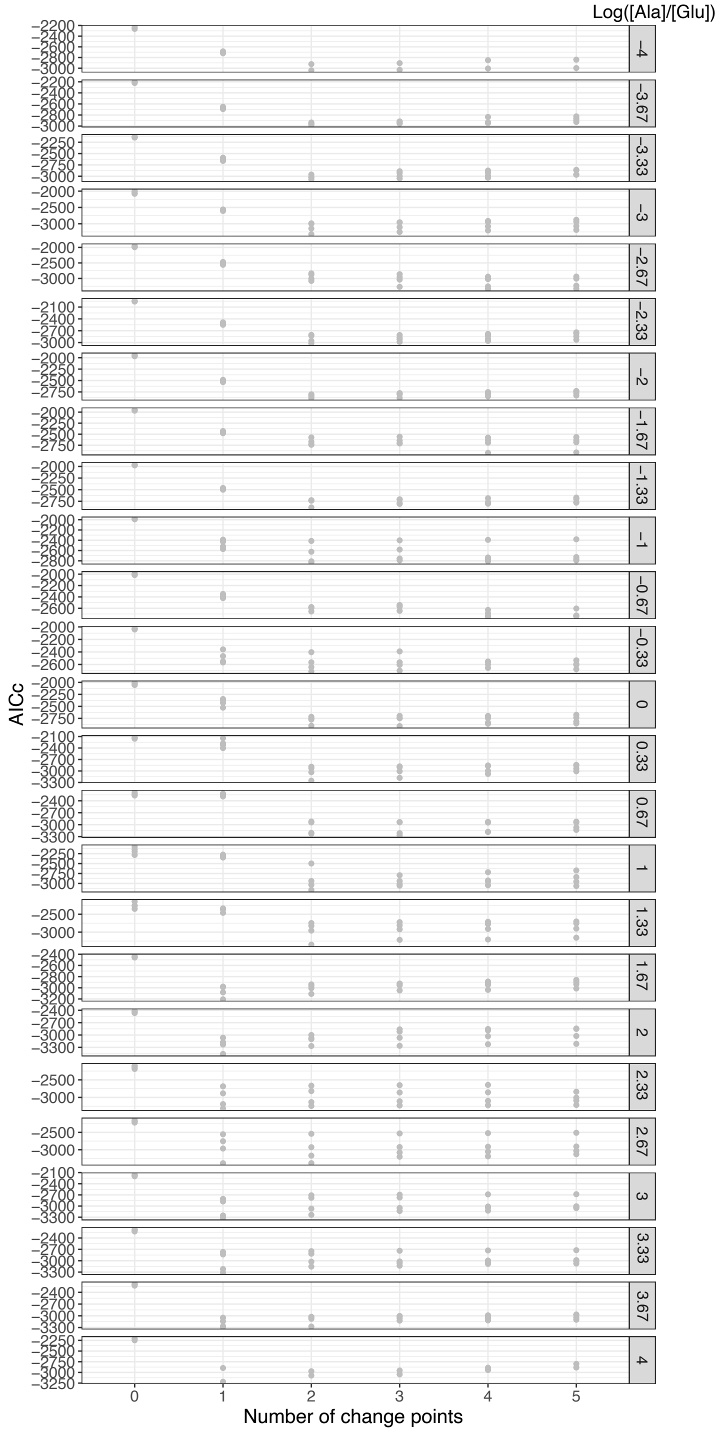


Supplementary Figure 5: **Segmented fits for the diauxic growth dataset:** AICc values for segmented fits of the dataset from Ref. [7] with varying numbers of change points.

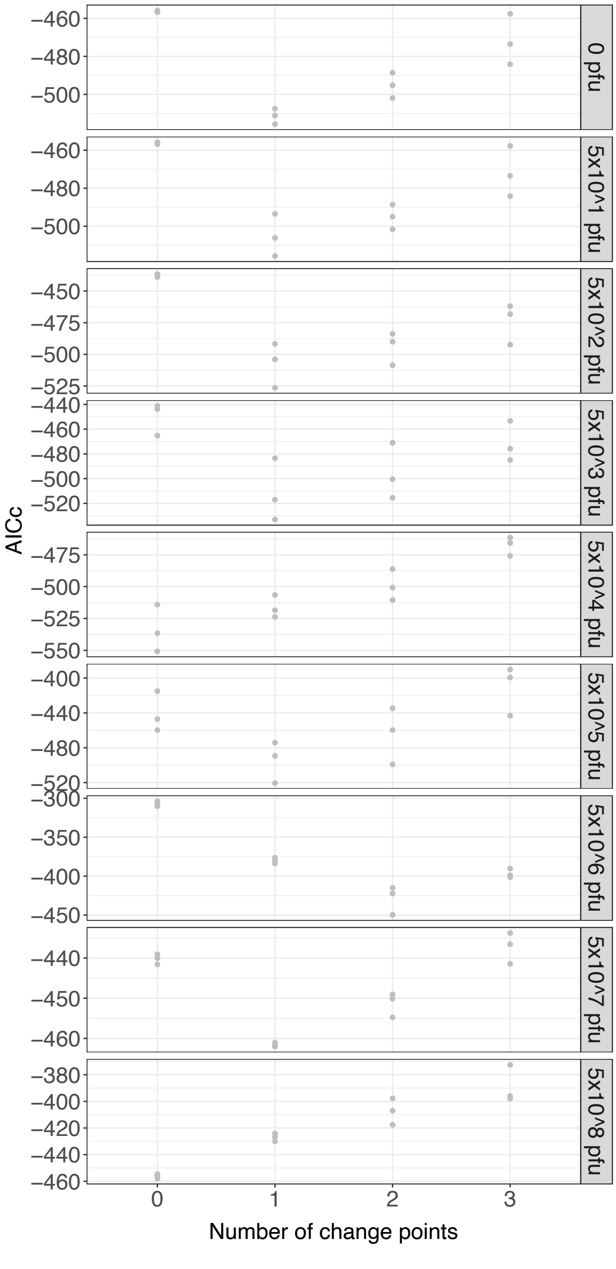


Supplementary Figure 6: **Segmented fits for the phage-bacteria interactions dataset:** AICc values for segmented fits of the dataset from Ref. [8] with varying numbers of change points.

### Supporting figures for the automatic inference of empirical laws

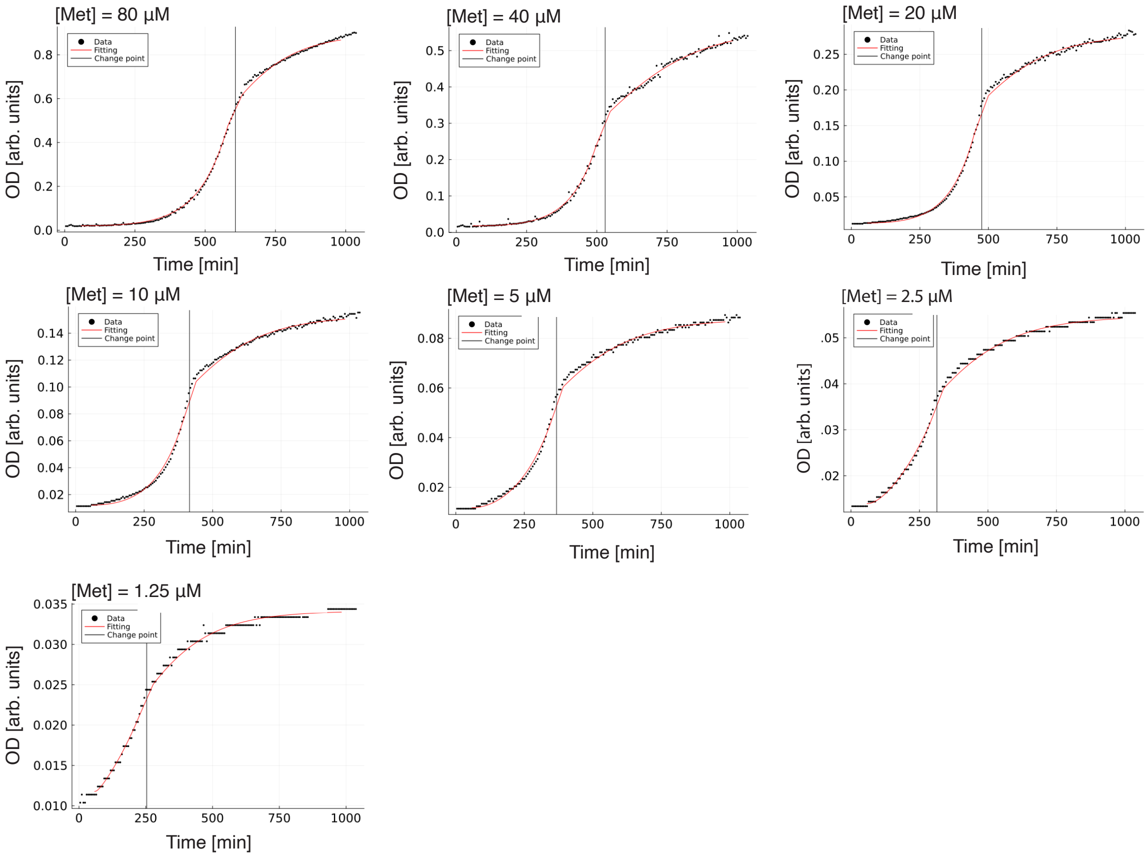


Supplementary Figure 7: **Model-based inference of auxotroph growth curves.** Measured growth curves of the $\Delta metA$ knockout (dots) together with segmented fit results (lines) for all growth experiments with supplemented methionine. Segmented fits were performed using one change point, with the exponential HPM (Eq. 25) and logistic models (Eq. 14) as candidates (ODE representation fits). The first segment was used to determine the exponential growth rate, while the second segment was used to estimate the saturation level or total growth.

**A**

| **Equation** | **Complexity** | **MSE** |
|---|---|---|
| $0.0089$ | 1 | $7.86 \times 10^{\ 6}$ |
| $([S] \times 5.57 \times 10^{\ 5}) + 0.0077$ | 5 | $5.70 \times 10^{\ 6}$ |
| $\frac{[S]}{[S]+1.35} \times 0.0114$ | 7 | $2.64 \times 10^{\ 7}$ |
| $\frac{0.0114}{([S]+0.18+1.13)} \times [S]$ | 9 | $2.61 \times 10^{\ 7}$ |
| $\left(\frac{0.0114}{(([S]^2+0.36\ \ 0.23)+[S])} \times [S]\right) \times [S]$ | 15 | $2.21 \times 10^{\ 7}$ |

**B**

| **Equation** | **Complexity** | **MSE** |
|---|---|---|
| $0.30$ | 1 | $0.11$ |
| $[S] \times 0.0132$ | 3 | $4.74 \times 10^{\ 4}$ |
| $([S] + 1.78) \times 0.0128$ | 5 | $1.64 \times 10^{\ 4}$ |
| $((([S] \times -2.76 \times 10^{\ 5}) + 0.0151) \times [S]$ | 7 | $1.04 \times 10^{\ 4}$ |
| $((([S] \times -2.76 \times 10^{\ 5}) + 0.0151) \times ([S] + 0.35)$ | 9 | $7.96 \times 10^{\ 5}$ |
| $(((([S] + 2.40) \times -2.76 \times 10^{\ 5}) + 0.0151) \times ([S] + 0.39)$ | 11 | $7.00 \times 10^{\ 5}$ |
| $(((([S] + ([S] \times -0.42)) \times -3.33 \times 10^{\ 5}) + 0.0143) \times ([S] + 0.86)$ | 13 | $4.52 \times 10^{\ 5}$ |

Supplementary Figure 8: **Automatic detection of empirical laws.** Hall of fame of symbolic regression for the growth rate (**A**) and total growth (**B**) for an *E. coli* $\Delta metA$ knockout expressing a red fluorescent marker (mCherry). Model candidates for equations representing the response of the observables as a function of substrate concentration $[S]$ are shown, along with the mean squared error and the complexity of the proposed models. As with the green-labeled $\Delta metA$ knockout discussed in the main text (cf. Fig. 4 and Extended Data Fig. 4), symbolic regression successfully recovers Monod's empirical laws (highlighted in the black boxes).

### Supporting figures for the characterization of chemical mixture landscapes

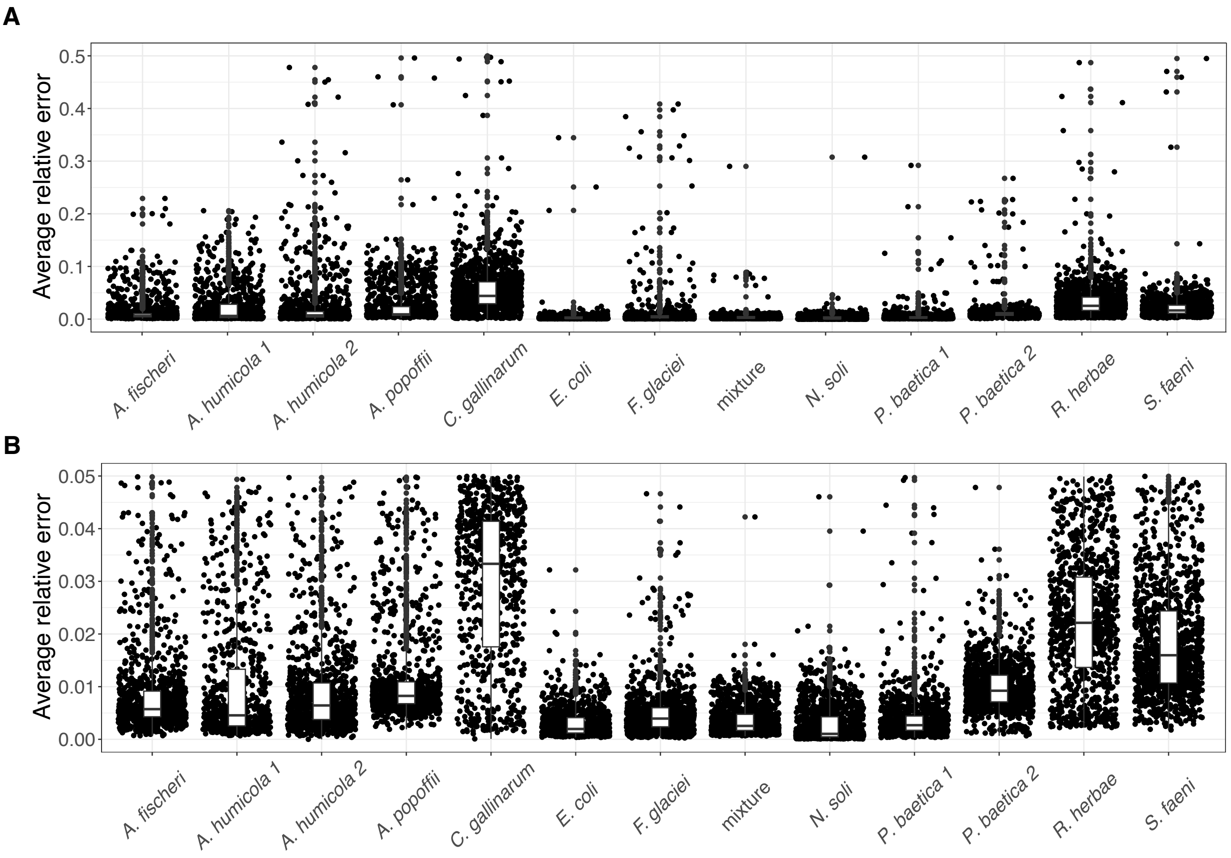


Supplementary Figure 9: **Average relative error for the fits of the dataset of Ref. [9].** **A**, Distributions of the average relative error by strain (number of samples: 15120). **B**, Distributions of the average relative error by strain after excluding misfits with an average relative error greater than 5% (number of samples: 13753). The fit results from these remaining samples were used in the analyses presented in the main text.

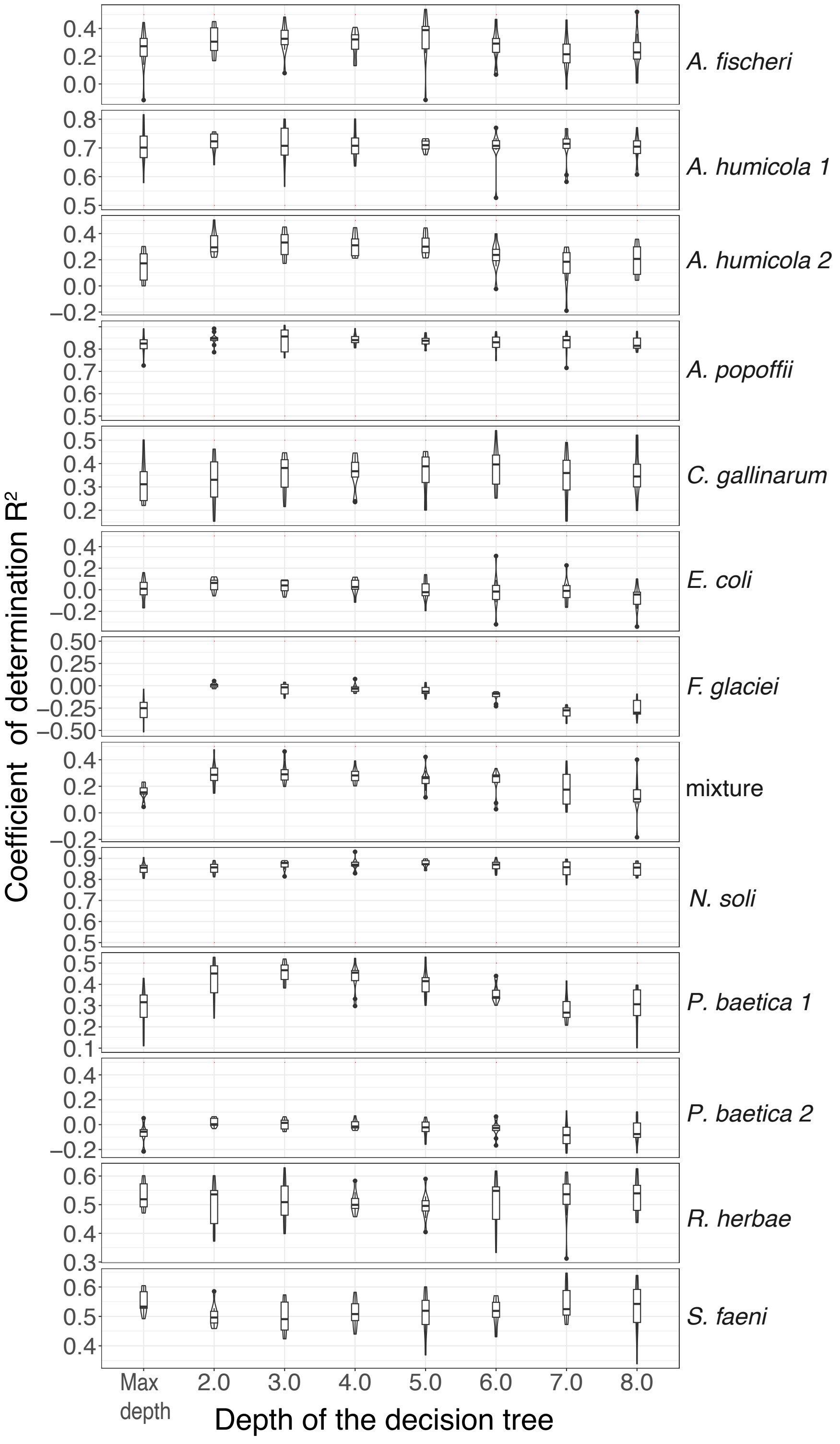


Supplementary Figure 10: **Decision tree cross validation for the growth rate**. Box and violin plots show the coefficients of determination $R^2$ for the 10-fold cross validation (10 runs) of decision tree applied to the growth rates inferred using the Richards model (Eq. 4) for the growth curves of all strains in the ecotoxicological dataset analyzed from Ref. [9].

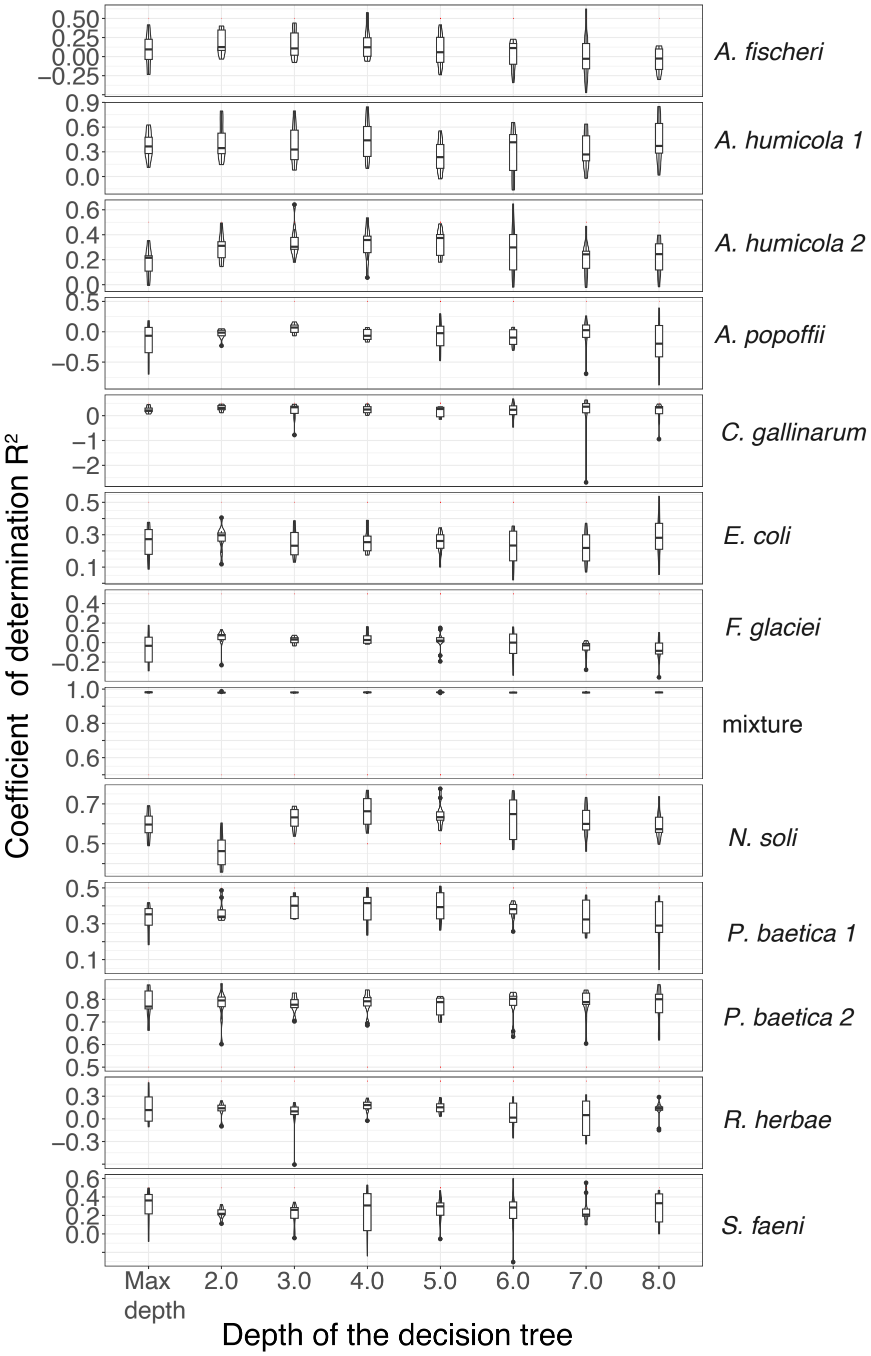


Supplementary Figure 11: **Decision tree cross validation for the total growth**. Box and violin plots show the coefficients of determination $R^2$ for the 10-fold cross validation (10 runs) of decision tree applied to the total growth inferred using the Richards model (Eq. 4) for the growth curves of all strains in the ecotoxicological dataset analyzed from Ref. [9].

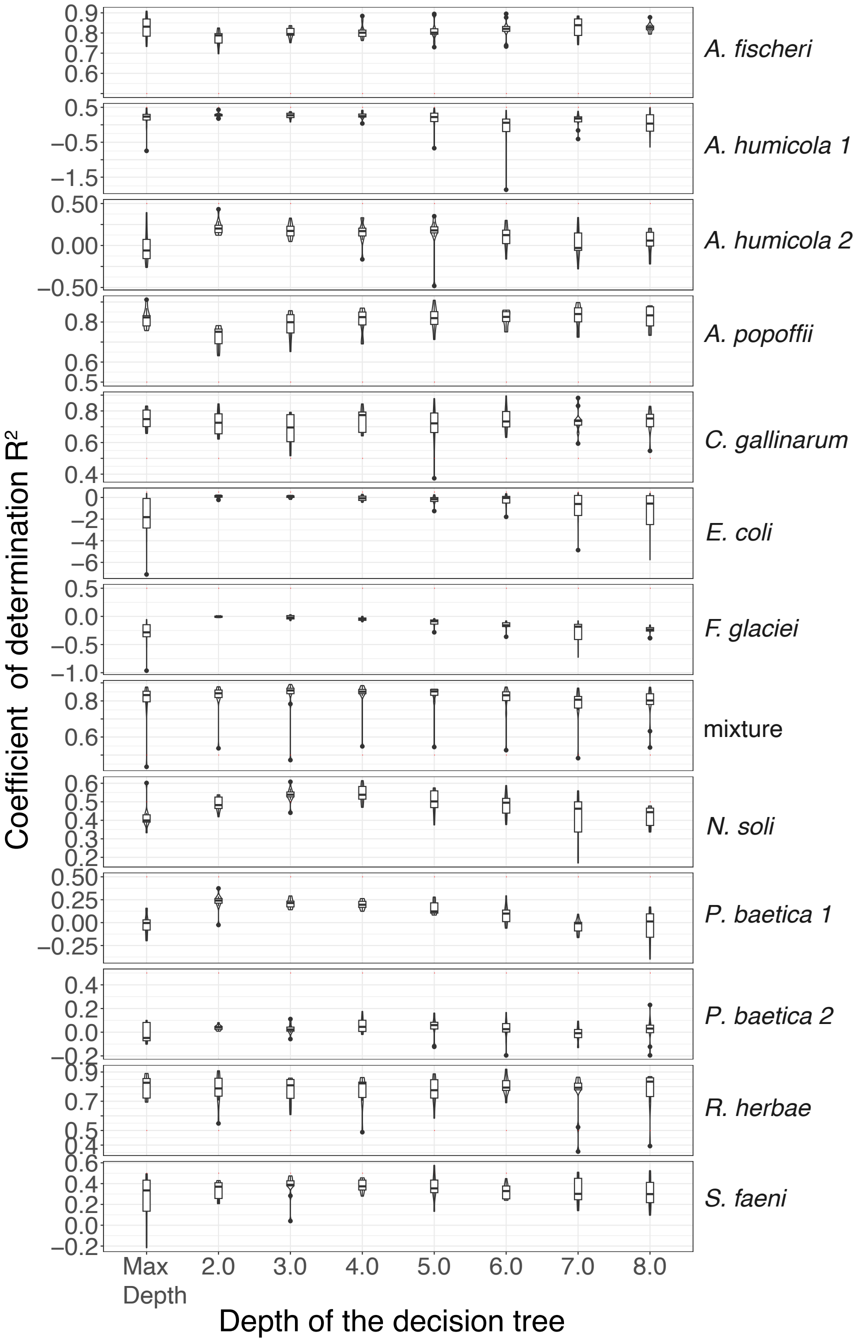


Supplementary Figure 12: **Decision tree cross validation for the lag time**. Box and violin plots show the coefficients of determination $R^2$ for the 10-fold cross validation (10 runs) of decision tree applied to the lag time inferred using the Richards model (Eq. 4) for the growth curves of all strains in the ecotoxicological dataset analyzed from Ref. [9].
